# Protein Corona Composition and Dynamics on Carbon Nanotubes in Blood Plasma and Cerebrospinal Fluid

**DOI:** 10.1101/2020.01.13.905356

**Authors:** Rebecca L. Pinals, Darwin Yang, Daniel J. Rosenberg, Tanya Chaudhary, Andrew R. Crothers, Anthony T. Iavarone, Michal Hammel, Markita P. Landry

## Abstract

When a nanoparticle enters a biological environment, the surface is rapidly coated with proteins to form a “protein corona”. Presence of the protein corona surrounding the nanoparticle has significant implications for applying nanotechnologies within biological systems, affecting outcomes such as biodistribution and toxicity. Herein, we measure protein corona formation on single-stranded DNA wrapped single-walled carbon nanotubes (ssDNA-SWCNTs), a high-aspect ratio nanoparticle ideal for sensing and delivery applications, and polystyrene nanoparticles, a model nanoparticle system. The protein corona of each nanoparticle is studied in human blood plasma and cerebrospinal fluid. We characterize corona composition by proteomic mass spectrometry to determine abundant and differentially enriched vs. depleted corona proteins. High-binding corona proteins on ssDNA-SWCNTs include proteins involved in lipid binding and transport (clusterin and apolipoprotein A-I), complement activation (complement C3), and blood coagulation (fibrinogen). Of note, albumin is the most common blood protein (55% w/v), yet exhibits low-binding affinity towards ssDNA-SWCNTs, displaying 1300-fold lower bound concentration relative to native plasma. We investigate the role of electrostatic and entropic interactions driving selective protein corona formation, and find that hydrophobic interactions drive inner corona formation, while shielding of electrostatic interactions allows for outer corona formation. Lastly, we study real-time binding of proteins on ssDNA-SWCNTs and find relative agreement between proteins that are enriched and bind strongly, such as fibrinogen, and proteins that are depleted and bind marginally, such as albumin. Interestingly, certain proteins express contrary behavior in single-protein experiments than within the whole biofluid, highlighting the importance of cooperative mechanisms driving selective corona adsorption on the SWCNT surface. Knowledge of the protein corona composition, dynamics, and structure informs translation of engineered nanoparticles from *in vitro* design to effective *in vivo* application.

## Introduction

Engineered nanoparticles are prominent in biological sensing, imaging, and delivery applications due to their distinctive optical and physical properties.^1^ The critical – yet often overlooked – challenge with these nanoscale tools is understanding the mechanisms of interaction between the nanoprobe and the biological system they are designed to query.^2,3^ Nanotechnologies are often developed, characterized, and validated *in vitro*, absent from the complexity of biological fluids.^4^ Yet when nanoparticles are introduced into a biological system, interfacially active proteins spontaneously adsorb to the foreign nanoparticle surfaces, leading to the formation of the “protein corona” ^5^ Binding of proteins to pristine nanoparticles can adversely affect the structure and function of the bound proteins,^6–8^ and carries the additional consequence of masking and re-defining the nanoparticle identity.^2,9,10^ The formed nanoparticle-corona complex, rather than the designed nanoparticle, then becomes the entity interacting with biological machinery.

Consequently, the *in vivo* trafficking, biodistribution, clearance, and biocompatibility of the nanoparticle become unpredictable.^2,9,11,12^ These corona-mediated alterations manifest as decreased nanoparticle efficacy or loss of *in vitro*-validated results, whereby the nanoparticle no longer carries out its designated function.^13,14^ Moreover, the protein corona is dynamic in nature, displaying a high degree of temporal heterogeneity.^2,15^ Rapid protein association and dissociation events on the nanoparticle surface, in conjunction with differential protein affinities in the corona, give rise to additional complications in understanding the timescales over which nanoparticles retain their pristine, corona-free, attributes within biological environments.

Corona composition and interactions driving formation as functions of nanoparticle type, biological environment, and time remain poorly understood.^16^ Prior work has investigated the role of nanoparticle surface charge, chemistry, size, and curvature as factors governing protein corona formation.^3,17–20^ Consequent work has sought to mitigate corona formation through nanoparticle surface passivation with polymers such as polyethylene glycol (PEG) to abrogate protein adsorption and sterically stabilize the nanoparticle, yet these strategies display variable efficacy.^21–23^ Although many studies classify protein corona composition around specific nanoparticle systems, there remains debate as to which protein and nanoparticle characteristics are most important in determining corona composition, and how different biological environments contribute to nanoparticle compositional and temporal corona heterogeneity.^3,24^ These studies are particularly difficult to perform with nanoparticles that are small, ~few nm, or of extreme aspect ratios, such as SWCNTs. Additionally, it remains status quo to test nanoparticle biofouling and biocompatibility in blood serum, a blood-based fluid rich in albumin, the most abundant blood plasma protein. The assumptions that serum is a representative biofluid for *in vivo* studies and that protein abundance in a native biofluid determines its relative abundance in a nanoparticle corona both stand to be refined.

Understanding protein corona formation is essential to design nanoparticles for *in vivo* applications that are robust and stable in biological environments. Our work focuses on single-walled carbon nanotubes (SWCNTs), a nanoparticle class that possesses unique optical and physical properties ideal for biological imaging, molecular sensing, and bio-delivery applications.^25–27^ SWCNTs are composed of a graphitic lattice structure in a high aspect ratio, cylindrical geometry, resulting in an intrinsic near-infrared (nIR) bandgap fluorescence over which biological samples are optically transparent,^28^ with no photobleaching threshold.^29^ This pristine carbon structure renders the surface hydrophobic, and SWCNTs accordingly exist as aggregated bundles in aqueous solution.^28^ To apply SWCNTs in aqueous biological systems, noncovalent functionalization with amphiphilic polymers imparts water solubility to the SWCNT while retaining the nIR-emissive electronic structure.^25^ Select polymers confer molecular recognition functionality when adsorbed to the SWCNT surface, such as the single-stranded DNA (ssDNA) sequences of (GT)_6_ and (GT)_15_ that are now commonly used to image neurotransmitter dopamine in the brain at spatiotemporal scales of relevance to endogenous dopaminergic neuromodulation.^27,30,31^ To design and apply these and other SWCNT-based nanoparticle technologies in biological systems, it is critical to understand the composition, dynamics, and dominant mechanisms of protein corona formation.

Herein, we explore protein corona formation, probed with a selective adsorption assay that is generalizable for different types of nanoparticles and biofluids. We focus on two nanoparticles: a model system of commonly studied polystyrene nanoparticles (PNPs)^4,6,15,17,32,33^ and an under-studied system of noncovalently functionalized SWCNTs. Protein corona formation on these nanoparticles was assessed in two biofluids of interest: blood plasma, a standard biofluid model system with high protein concentration, relevant for blood circulation applications, and cerebrospinal fluid (CSF), an under-studied biological environment with low protein concentration, relevant for brain and other central nervous system studies. To the best of our knowledge, this is the first time that the corona formed on SWCNTs in CSF has been studied, and a knowledge of this corona is imperative for developing SWCNT-based applications in the brain. Corona composition was characterized by gel electrophoresis and quantitative, label-free mass spectrometry (MS) analysis, revealing key protein corona contributors and isolation of protein factors governing corona formation: functional roles in lipid binding and complement activation, and high content of certain charged and aromatic amino acids. By varying incubation conditions (dynamics, temperature, ionic strength), we identify factors governing protein adsorption. We posit that hydrophobic interactions dominate the formation of the “hard” inner corona, in which proteins interact directly with the nanoparticle surface, while electrostatic interactions govern the formation of the “soft” outer corona. Protein corona binding thermodynamics and kinetics were next assessed to measure adsorption of key proteins to (GT)_15_-SWCNTs via isothermal titration calorimetry (ITC) and a corona exchange assay.^34^ This quantification of the time-dependent protein corona provides a comprehensive understanding of the interactions driving the corona formation process. Finally, protein corona structure was visualized by small angle x-ray scattering (SAXS) and transmission electron microscopy (TEM). SAXS reveals changing mass fractal morphology of the ssDNA-SWCNTs in the presence of a high-binding protein (fibrinogen) that is otherwise absent in the case of the low-binding protein (albumin), and gives rise to a physical picture of the final protein-SWCNT complex structure and higher-order aggregate formation.

As nanoparticles are increasingly implemented as tools to study and alter biological systems, it is crucial to develop an understanding of how these nanoparticles interact with biosystems. In this work, we conduct a multimodal study on the mechanisms governing the composition, driving forces of formation, and binding process of protein-nanoparticle complexes. This understanding will in turn serve to establish parameters for the rational design of future nanobiotechnologies and to allow more reliable translation from *in vitro* design to *in vivo* applications in protein-rich environments, including the brain.

## Results and Discussion

### Validation of modified pull-down assay to study protein corona composition

We employed a modified pull-down assay (**Figure 1**) to investigate the end-point corona composition (see **Modified pull-down assay** in Materials and Methods and further details in SI).^17^ With this assay, selective adsorption of proteins onto nanoparticles was evaluated by (i) incubating nanoparticles with the target biological fluid for 1 h, (ii) pulling down protein-nanoparticle complexes by centrifugation, (iii) removing unbound proteins remaining in solution with three washing steps, (iv) eluting selectively bound proteins from the nanoparticle surface with buffer containing surfactant and reducing agent, and (v) characterizing proteins by two-dimensional polyacrylamide gel electrophoretic separation (2D PAGE) or liquid chromatography-tandem mass spectrometry (LC-MS/MS). Each step of the assay was validated by confirming proteins adsorb to and desorb from polystyrene nanoparticle (PNP) surfaces as predicted during the workflow: dynamic light scattering (DLS) tracked the increase in protein-PNP hydrodynamic radius, solution absorbance confirmed protein-PNP association followed by removal of unbound protein, and quantifying eluted protein verified full desorption of the PNP protein corona prior to composition characterization (**Figure S1**). Furthermore, zeta potential measurements of blood plasma proteins alone, nanoparticles alone, and protein-nanoparticle conjugates (**Figure S2**) reveal that both separate entities were initially negatively charged in buffered solution, whereby the formation of the protein corona on the nanoparticles reduced the magnitude of the zeta potential. The measured reduction in effective surface charge of the conjugates confirms the colloidal instability observed upon combining nanoparticles with plasma, in agreement with previous literature.^4^

**Figure 1.**
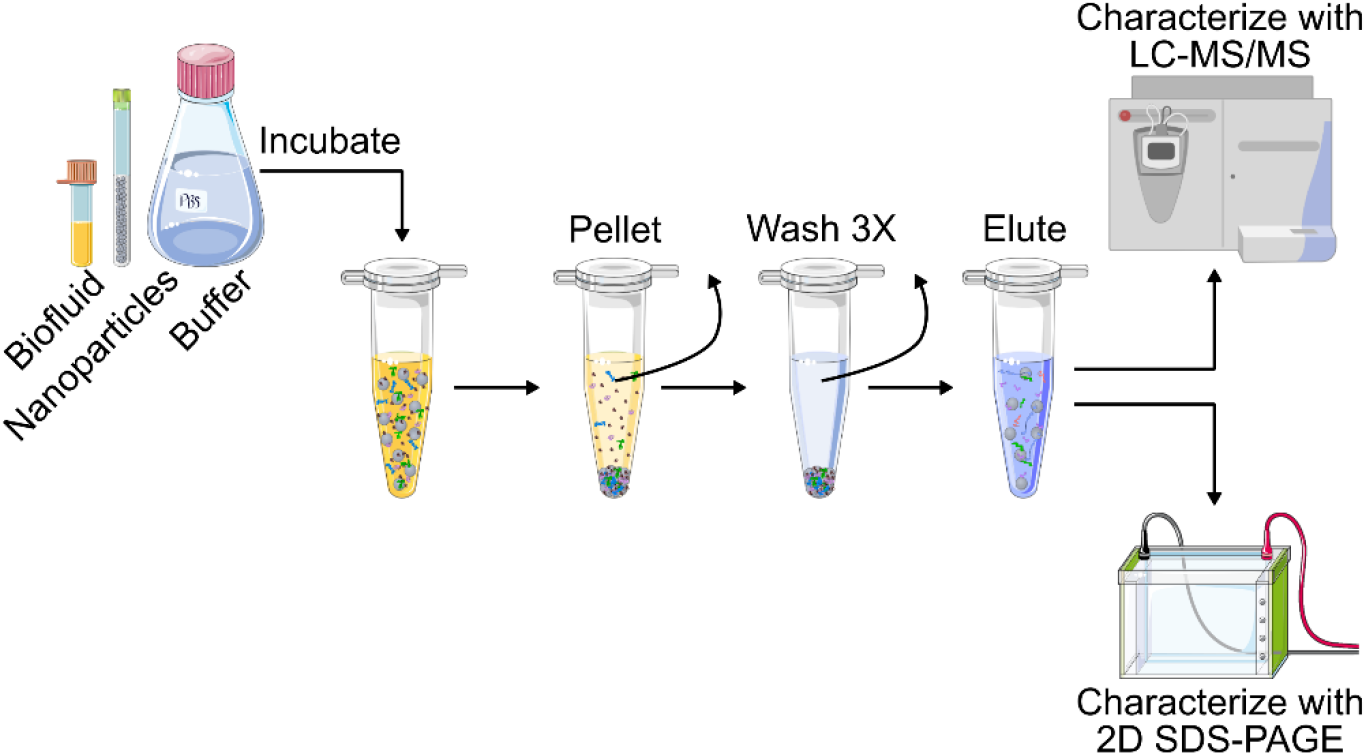
Modified pull-down assay to determine protein corona composition on nanoparticles. Schematic detailing experimental procedure to study protein corona formation. Nanoparticles are incubated with the desired biofluid in buffered solution, the nanoparticle-protein complexes are pelleted by centrifugation and washed three times to remove non-selectively pulled-down proteins, and corona proteins are eluted and characterized by two-dimensional polyacrylamide gel electrophoresis (2D PAGE) or liquid chromatography-tandem mass spectrometry (LC-MS/MS).

### Nanoparticle protein corona composition overview

Following validation of our modified pull-down assay, the spontaneous formation of nanoparticle coronas was studied on two distinct nanoparticle surfaces: PNPs and (GT)_15_-SWCNTs. Two different biofluid exposures were tested: human blood plasma and human cerebrospinal fluid. Following the modified pull-down workflow depicted in **Figure 1**, each nanoparticle was incubated with either biofluid prior to eluting corona proteins for identification by 2D PAGE. PAGE analysis confirmed the presence of abundant proteins in native blood plasma and CSF, and eluted proteins from nanoparticles showed selective enrichment or depletion fingerprints (**Figure S3**). More in-depth protein corona composition studies were subsequently undertaken by performing in-solution trypsin digestion of proteins eluted from nanoparticles, followed by protein characterization with label-free, quantitative mass spectrometry (LC-MS/MS). Analysis by LC-MS/MS provided information on: (i) molar corona protein abundances via comparison to an internal standard (see **LC-MS/MS** in Materials and Methods) and (ii) enrichment or depletion in each nanoparticle corona, relative to protein concentrations in the native biofluid. To assess enrichment or depletion, we define fold change as the ratio of molar corona protein abundance to molar biofluid protein abundance, normalized by total protein mass in the respective samples (see further explanation in SI). The twenty most abundant proteins in the nanoparticle coronas are summarized in **Table 1** for plasma and **Table 2** for CSF. The highest concentration proteins in the native biofluids are both in relative agreement with previous literature (with the exception of fibrinogen, as detailed in SI).^35–38^ Proteins identified in the corona formed on PNPs incubated in plasma are corroborated by previous literature,^15,17^ further validating our pull-down assay. Protein abundance and fold-change (relative to the native biofluid alone) for protein coronas formed on PNPs and (GT)_15_-SWCNTs are represented graphically in **Figure 2** for incubation in blood plasma and **Figure 4** for incubation in CSF. Here, proteins are grouped by functional class according to PANTHER.^39^ Full protein lists are included in SI (**Figure S4** for plasma, **Figure S5** for CSF, and tabulated in attached file).

**Table 1.** Top 20 most abundant proteins identified by proteomic mass spectrometry in plasma alone and in each nanoparticle corona.

|  | Plasma alone | PNPs + plasma | (GT)<sub>15</sub>-SWCNTs + plasma |
| --- | --- | --- | --- |
| 1 | Serum albumin | Alpha-2-HS-glycoprotein | Clusterin |
| 2 | Haptoglobin | Ig kappa constant | Histidine-rich glycoprotein |
| 3 | Ig kappa constant | Haptoglobin | Apolipoprotein A-I |
| 4 | Ig heavy constant gamma | Complement C3 | Complement C3 |
| 5 | Serotransferrin | Kininogen-1 | Haptoglobin |
| 6 | Apolipoprotein A-I | Ig heavy constant gamma 1 | A disintegrin and metalloproteinase with thrombospondin motifs 12 |
| 7 | Complement C4 | Apolipoprotein A-II | Complement C1r subcomponent |
| 8 | Telomeric repeat-binding factor 2-interacting protein | tRNA-dihydrouridine(47) synthase [NAD(P)(+)]-like | Vitronectin |
| 9 | Alpha-1-antitrypsin | Beta-2-glycoprotein 1 | Kininogen-1 |
| 10 | Alpha-2-HS-glycoprotein | Vitronectin | Prothrombin |
| 11 | Apolipoprotein A-II | Serum albumin | C4b-binding protein alpha chain |
| 12 | Ig heavy constant alpha 1 | Vitamin D-binding protein | Complement factor H |
| 13 | Integrin alpha-7 | A disintegrin and metalloproteinase with thrombospondin motifs 12 | Fibrinogen alpha chain |
| 14 | Alpha-2-macroglobulin | Hemopexin | Protein AMBP |
| 15 | Complement C3 | Apolipoprotein A-I | Beta-2-glycoprotein 1 |
| 16 | Complement C5 | Ig lambda-like polypeptide 5 | Apolipoprotein E |
| 17 | Hemopexin | Histidine-rich glycoprotein | Complement C1q subcomponent subunit B |
| 18 | Alpha-1-acid glycoprotein 1 | Clusterin | Ig heavy constant gamma 1 |
| 19 | Ig heavy constant mu | Alpha-1-antitrypsin | Ig J chain |
| 20 | Beta-2-glycoprotein 1 | Serum paraoxonase/arylesterase 1 | Galectin-3-binding protein |

**Table 2.**
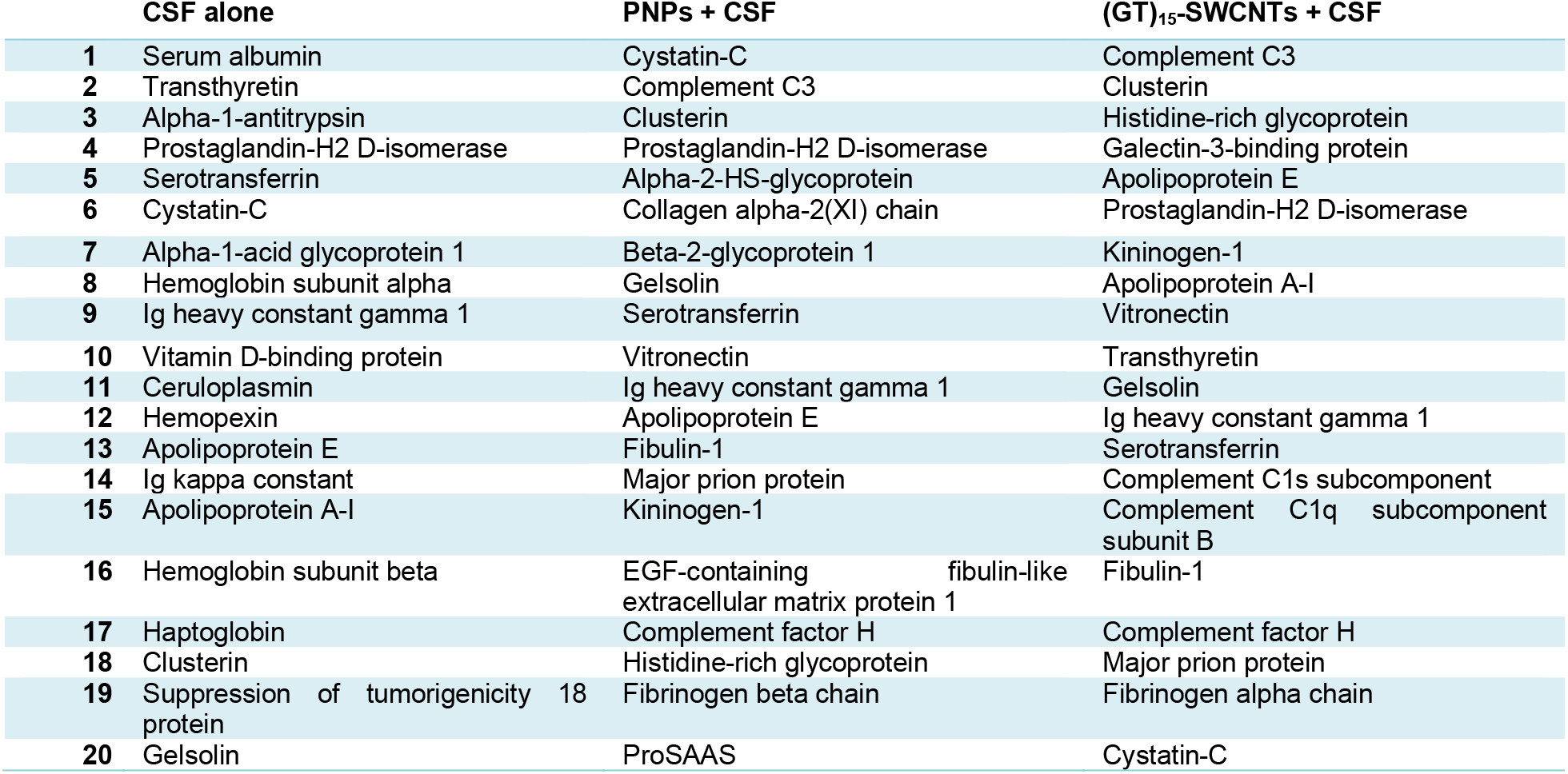
Top 20 most abundant proteins identified by proteomic mass spectrometry in CSF alone and in each nanoparticle corona.

**Figure 2.**
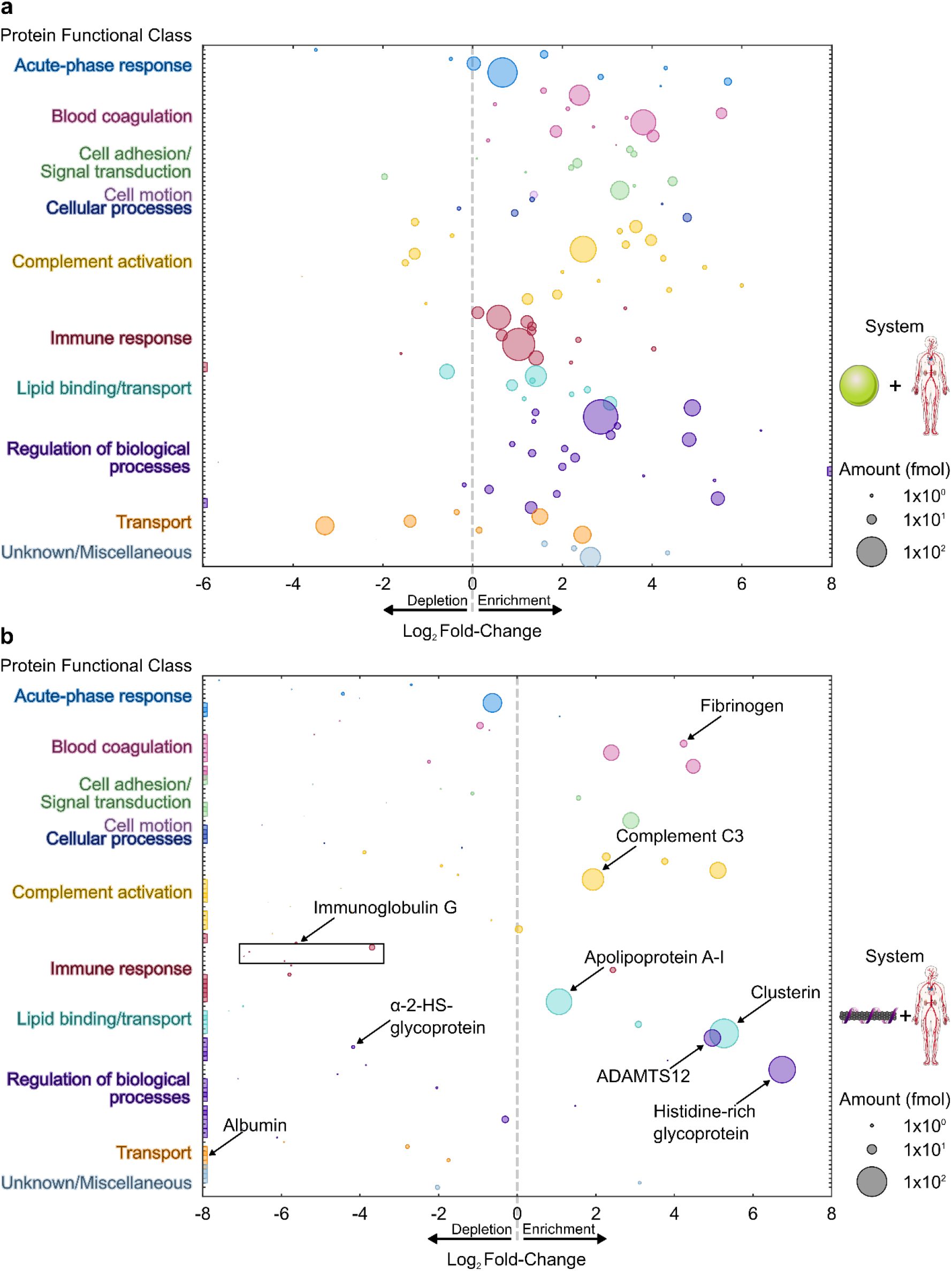
Blood plasma protein corona composition determined by proteomic mass spectrometry. Protein corona formed from blood plasma on **(a)** PNPs and **(b)** (GT)_15_-SWCNTs. Proteins are grouped by functional class according to PANTHER.^39^ Full protein lists are available in SI (Figure S4 and attached tables). Log_2_ fold-change is in comparison to the native biofluid alone. Circle size corresponds to the protein abundance (femtomolar). Colored boxes at x-axis limits indicate no protein detected in either corona (x < 2^−6^ or 2^−8^) or biofluid (x > 2^8^).

### Blood plasma protein corona composition and functional roles

Our LC-MS/MS analysis highlights the significant enrichment vs. depletion of specific plasma proteins in the (GT)_15_-SWCNT plasma corona. Both LC-MS/MS and 2D PAGE show that the most abundant protein in plasma, serum albumin (55% w/v in plasma), does not appreciably interact with (GT)_15_-SWCNTs: albumin is in low abundance in the protein corona and significantly depleted, with a 0.00077-fold change relative to native plasma albumin composition. The most highly enriched protein, histidine-rich glycoprotein (107-fold enrichment), is hypothesized to interact with other plasma proteins to enter the corona.^4^ Another greatly enriched protein, unreported in prior carbon nanoparticle corona literature, is ‘a disintegrin and metalloproteinase with thrombospondin motifs 12’ (ADAMTS12), which appeared in high abundance on both (GT)_15_-SWCNTs (6^th^ highest abundance) and (GT)_6_-SWNCTs (1^st^ highest abundance). ADAMTS12 is a metalloprotease with a zinc cofactor thought to possess anti-tumorigenic properties and to play a role in cell adhesion, pointing to potential applications in protein-SWCNT construct design.

For both PNP and (GT)_15_-SWCNT nanoparticles in plasma, the protein corona displayed a distribution of enriched, highly abundant proteins across different functional roles (**Figure 2**). Corona proteins on PNPs are more diverse in protein functional classes than corona proteins on (GT)_15_-SWCNTs, representing a range of endogenous functions including adaptive immune response and transport; functions for which proteins are largely absent in the plasma corona of (GT)_15_-SWCNTs. To quantify the broad functional roles of plasma proteins involved in corona formation, we regressed ln-fold change against protein class using effect-coding while controlling for sample-to-sample variability (see **Linear regression models** in Materials and Methods). The regression coefficient for each protein class in each biofluid-nanoparticle system (first two columns of **Figure 3**) quantifies the fractional fold change of that protein class relative to the average fold change of all protein classes. On PNPs in plasma, no class of protein is substantially different from the average fold change, whereas on (GT)_15_-SWCNTs, several protein classes diverge from the average fold change. These dissimilarities highlight the more agnostic association of proteins to the PNP surface in comparison to the (GT)_15_-SWCNT surface, where this improved ability of (GT)_15_-SWCNTs to resist nonspecific protein biofouling presents a promising outcome towards SWCNT-based biotechnology applications.

**Figure 3.**
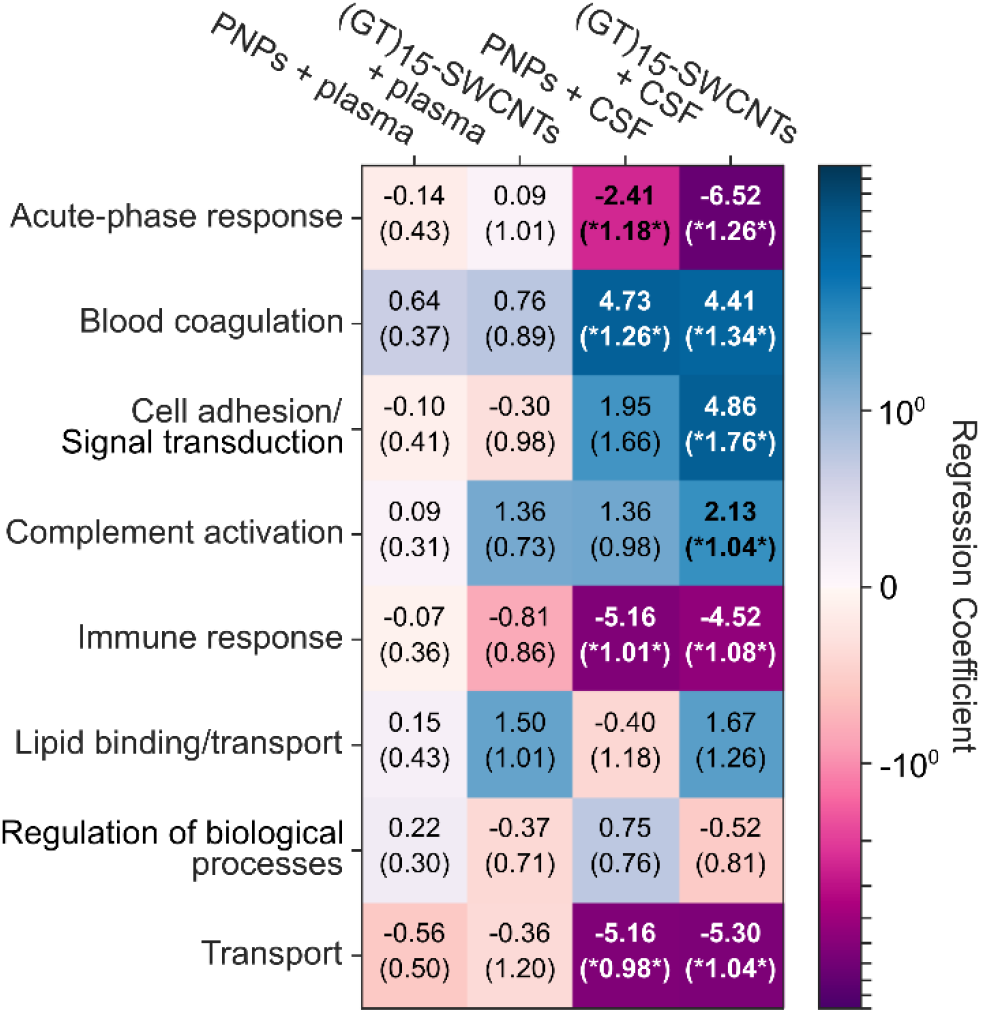
Role of protein functional class in protein corona formation for each nanoparticle-biofluid pairing. Ln-fold change, effect-coded regression coefficients of protein classes (rows) for each nanoparticle-biofluid pairing (columns). Cells are colored from dark purple (lower than the average fold change) to white (average fold change) to dark blue (higher than average fold change). Standard errors of the coefficients are given in parentheses. Results that have false-discovery-rate-corrected p-values of below 0.1 are bolded and noted with asterisks.

More specifically, plasma proteins enhanced on (GT)_15_-SWCNTs are involved in (i) lipid binding/transport (150% fold change over the average of all protein classes) and (ii) complement activation (140% fold change) (**Figure 3**). A key example of a lipid binding/transport corona protein is clusterin (a.k.a. apolipoprotein J) as the most abundant plasma protein in the (GT)_15_-SWCNT corona (38-fold enrichment; **Figure 2**), adsorption of which has shown promise in reducing non-specific cellular uptake of other types of nanocarriers.^32^ Apolipoproteins broadly act as dysopsonins that promote prolonged circulation in the blood.^4^ Hydrophobic interactions are posited to drive apolipoprotein adsorption in mimicry of native functions, such as apolipoprotein A-I (3^rd^ most abundant) that binds and transports hydrophobic fats through aqueous environments.^15,40^ We expect apolipoprotein binding, including clusterin and apolipoprotein A-I, to have a considerable impact on intracellular trafficking and fate of nanoparticles *in vivo*.^17,41^ Regarding group (ii), complement C3 is the 4^th^ most abundant plasma protein on (GT)_15_-SWCNTs, with a 14-fold enrichment. This result is in agreement with previous literature that SWCNTs activate complement via the classical pathway.^40,42^ Binding of complement proteins leads to nanoparticle opsonization if the bound proteins are intact, which could be useful in developing targeted therapies, yet detrimental if longer bloodstream circulation time is desired.^3,4^ In contrast to the high representation of complement proteins in the corona that serve a role in the innate immune response, it is interesting to note the low corona representation of immunoglobulins (81% lower fold change than average, **Figure 3**), proteins involved in the adaptive immune response. In addition to groups (i) and (ii), blood coagulation proteins are enhanced 76% on (GT)_15_-SWNCTs, but the wide distribution of regression coefficients for these proteins precludes making statistically significant conclusions about this class (**Figure S7**). A notable enriched blood coagulation protein is fibrinogen, with19-fold enrichment in the (GT)_15_-SWCNT corona relative to concentrations in plasma. Fibrinogen’s presence in the corona is unfavorable, as fibrinogen is responsible for eliciting inflammatory responses and nanoparticle aggregation.^4,8,43^

Our enrichment and depletion results for (GT)_15_-SWCNTs in blood plasma have practical implications for nanoparticle studies *in vivo*. Specifically, albumin composes 55% (w/v) of proteins in blood plasma, corresponding to 35-50 mg/mL,^38^ and is often assumed to comprise a representative constituent in nanoparticle protein coronas. In consequence, many nanotechnologies are tested for functionality in serum instead of plasma.^1,47,48^ Although albumin alone has been known to disperse SWCNTs in aqueous solution under sonication conditions,^12,46^ here we note that albumin is likely unable to outcompete higher affinity proteins when in the presence of a complex biofluid. Accordingly, we hypothesize that albumin plays a minimal role in the full plasma protein corona and subsequent *in vivo* trafficking and fate. This finding of low albumin adsorption is important for redefining interactions of ssDNA-SWCNT complexes with *in vivo* systems, and the standards by which nanotechnologies are tested are based on these interactions.

To test consistency in protein corona formation on SWCNTs, we further compared the plasma corona formed around (GT)_15_-SWCNTs to that on (GT)_6_-SWCNTs, where the ssDNA strand adsorbed to the SWCNT differs in length (30 vs. 12 nucleotides, respectively).^30^ The plasma proteins identified in the (GT)_6_-SWCNT corona approximately match those in the (GT)_15_-SWCNT corona (**Figure S4** and **Table S1**), with 85% match within the top 20 most abundant proteins.

### Cerebrospinal fluid protein corona composition and functional roles

We repeated our assay and analysis to study protein corona formation from CSF on both PNPs and (GT)_15_-SWCNTs. Highly abundant proteins in the CSF corona formed on (GT)_15_-SWCNTs include complement C3 as the most abundant (26-fold enriched relative to native CSF), clusterin as the 2^nd^ most abundant (15-fold enriched), and histidine-rich glycoprotein as the 3^rd^ most abundant (41-fold enriched). Again, serum albumin is remarkably absent, with a 3.9×10^−7^-fold depletion in comparison to CSF. Notably, a reproducible outlier emerges from LC-MS/MS studies of (GT)_15_-SWCNTs incubated with CSF: galectin-3-binding protein (G3BP), which is the most strongly enriched (80-fold) on the SWCNT surface relative to native G3BP concentrations in CSF (relatively low in healthy individuals) and the fourth most abundant corona protein. This high degree of adsorption and enrichment of G3BP on (GT)_15_-SWCNTs is relative to not only CSF alone, but also to the corona formed on PNPs. Identification of highly adsorbing and potentially interfering proteins could enable *a priori* design of future nanosensors to either promote selective protein adsorption or mitigate unfavorable protein adsorption.

LC-MS/MS analysis of the CSF-based protein corona also revealed proteins across a range of functional classes for both nanoparticles (**Figure 3** and **Figure 4**). Many of the same key proteins from plasma are enriched when nanoparticle coronas are formed instead in CSF. Broad protein classes that have higher than average fold change on (GT)_15_-SWCNTs are blood coagulation proteins (441% higher fold change than average), complement proteins (213% higher), and cell adhesion/signal transduction proteins (486% higher). Protein classes substantially lower than average in the corona consist of acute-phase response proteins (653% lower), immune response (452% lower), and transport proteins (530% lower).

**Figure 4.**
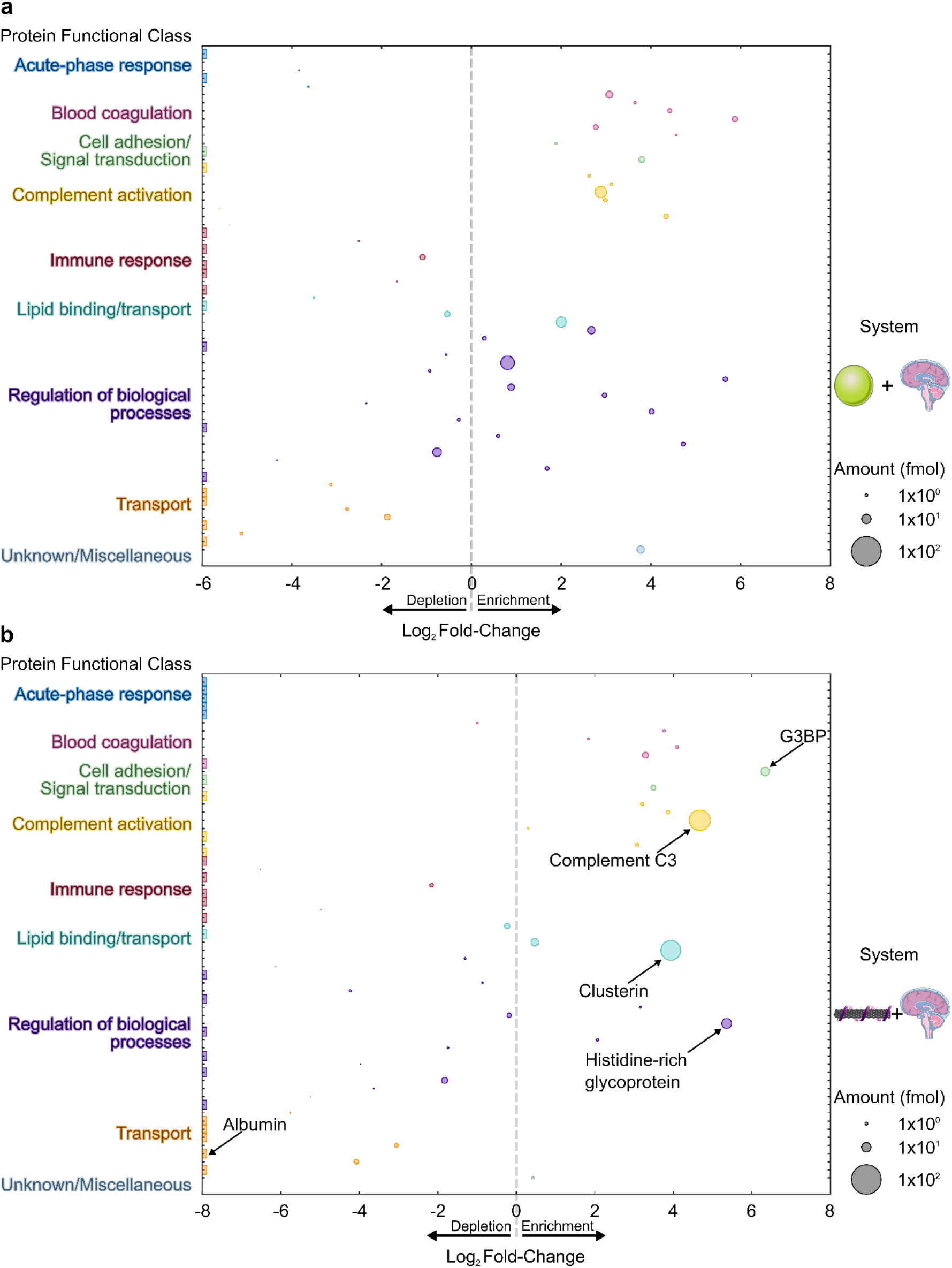
Cerebrospinal fluid (CSF) protein corona composition determined by proteomic mass spectrometry. Protein corona formed from CSF on **(a)** PNPs and **(b)** (GT)_15_-SWCNTs. Proteins are grouped by functional class according to PANTHER.^39^ Full protein lists are available in SI (**Figure S5** and attached tables). Log_2_ fold-change is in comparison to the native biofluid alone. Circle size corresponds to the protein abundance in femtomolar quantities. Colored boxes at x-axis limits indicate no protein detected in either corona (x < 2^−6^ or 2^−8^) or biofluid (x > 2^8^).

### Comparisons of plasma vs. cerebrospinal fluid protein corona compositions on (GT)_15_-SWCNTs

Many proteins have the same corona representation on (GT)_15_-SWCNTs across plasma and CSF. Examples of these conserved, highly abundant proteins on (GT)_15_-SWCNTs include clusterin, histidine-rich glycoprotein, and complement C3. However, certain proteins show distinctive behaviors in the (GT)_15_-SWCNT corona formed from different biofluids, such as G3BP increasing from #20 in plasma to #4 in CSF, despite G3BP’s higher native concentration in plasma than CSF, or serotransferrin missing from the plasma corona and present in the CSF corona, despite serotransferrin’s higher native concentration in plasma than CSF. These discrepancies point to complex mechanisms such as the Vroman effect,^47^ whereby surface adsorption is dictated by relative affinities and abundances of all protein constituents in the bulk phase to determine the end-state corona. Specifically, the Vroman effect predicts that highly abundant, lower affinity proteins will be replaced over time with less abundant, higher affinity proteins, a phenomenon that depends on minute differences in the protein environment. We also find that while plasma protein content in the corona vs. in the native biofluid is positively correlated for plasma proteins on PNPs (R^2^ = 0.461), this scaling does not hold for either (GT)_15_-SWCNTs (R^2^ = 0.101) or (GT)_6_-SWCNTs (R^2^= 0.072) (**Figure S6a-c**). Interestingly, corona abundance exhibits a weakly negative correlation with native abundance for CSF proteins on both PNPs (R^2^ = 0.012) and (GT)_15_-SWCNTs (R^2^ = 0.076) (**Figure S6d,e**). This again suggests complex mechanisms driving selective corona adsorption on SWCNTs and, depending on the biofluid, also PNPs.

### Molecular phenomena involved in protein corona formation

To evaluate the nanoscale mechanisms involved in corona formation, we linearly regressed the ln-fold change of proteins in the corona relative to the native biofluid against protein physicochemical properties including mass, post-translational modification frequency, binding site frequency, and amino acid percent compositions (see **Linear regression models** in Materials and Methods).^48,49^ Statistically, the calculated regression coefficients (**Figure 5** and distributions in **Figure S8**) quantify the fractional difference of the fold change for a protein with a unit increase of the independent variable, holding all other independent variables constant. Thermodynamically, the regression coefficients quantify the free energy change of a protein adsorbing into the corona per unit of the independent variable in units of k_b_T (see full derivation in SI).

**Figure 5.**
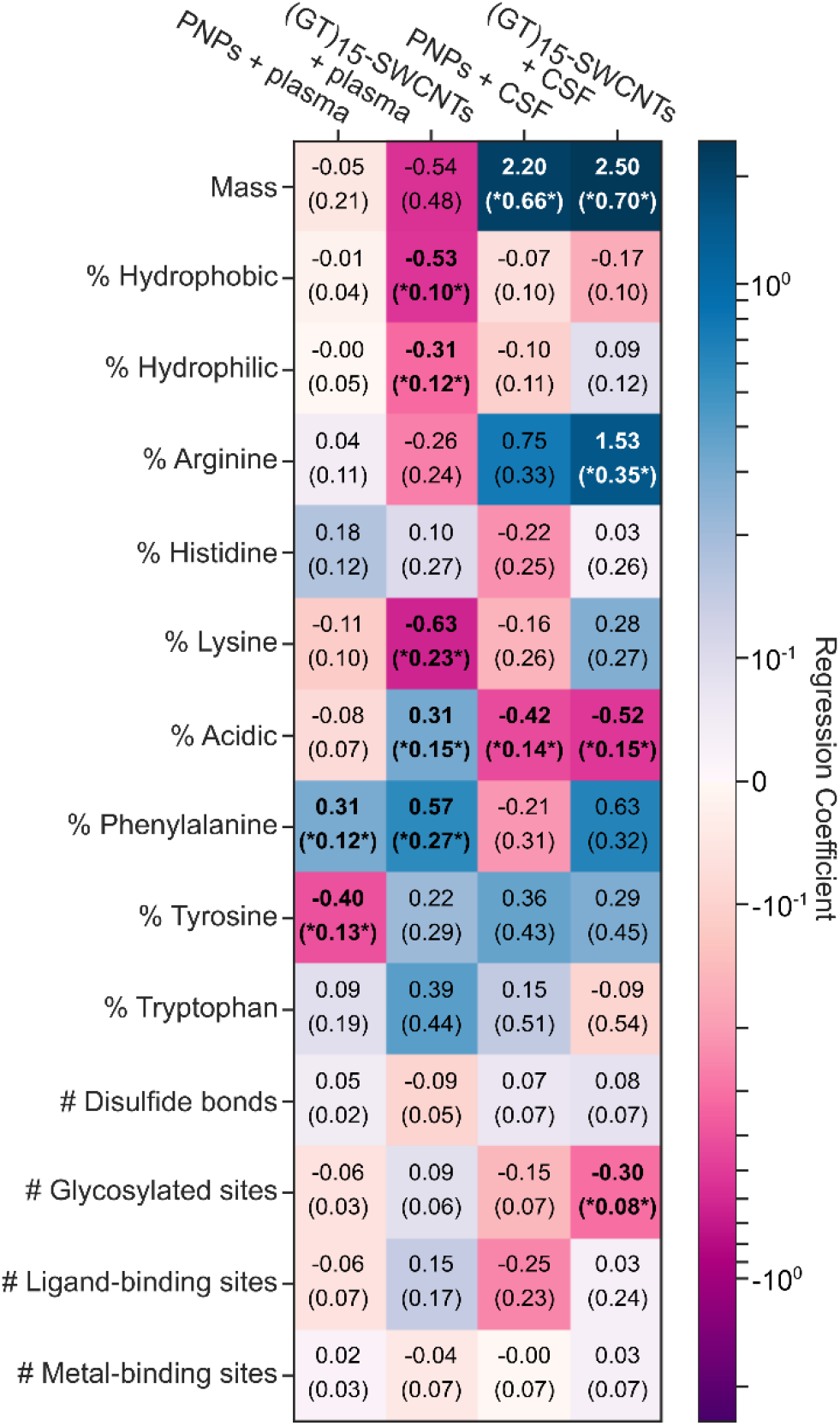
Molecular attributes of proteins that govern protein corona formation for each nanoparticle-biofluid pairing. Ln-fold change regression coefficients for molecular attributes of proteins (rows) for each nanoparticle-biofluid pairing (columns). Cells are colored from dark purple (negative effect on fold change) to white (no effect) to dark blue (positive effect). Standard errors of the coefficients are given in parenthesis. Results that have false-discovery-rate-corrected p-values of below 0.1 are bolded and noted with asterisks. Amino acid groupings include: non-aromatic hydrophobic (sum of alanine, valine, isoleucine, leucine, and methionine content), hydrophilic (sum of serine, threonine, asparagine, glutamine content), and acidic (sum of aspartic acid and glutamic acid content).

The plasma corona formed on PNPs is not possible to differentiate by protein molecular properties, except for a slight favorable interaction associated with the aromatic phenylalanine residue and an unfavorable interaction with tyrosine. Together with our protein abundance data, these results suggest protein corona formation on PNPs is diverse with respect to protein functional class and not selective for specific protein properties. In contrast, the plasma corona formed on (GT)_15_-SWCNTs is more selective, showing unfavorable interactions with non-aromatic, hydrophobic amino acids (−0.53 k_b_T / %hydrophobic) as well as polar amino acids (−0.31 k_b_T / %polar). The former result is surprising in that the SWCNT surface is extremely hydrophobic yet does not interact favorably with aliphatic residues. However, we find aromatic residues are enhanced in the (GT)_15_-SWCNT plasma corona, namely phenylalanine (0.57 k_b_T / %F), tyrosine (0.22 k_b_T / %Y), and tryptophan (0.39 k_b_T / %W). Regarding basic residue content, lysine (and to a lesser extent, arginine) is also associated with unfavorable interactions with (GT)_15_-SWCNTs (−0.63 k_b_T / %K), which is surprising in that positively charged proteins are expected to have favorable electrostatic interactions with the negatively charged, solvent-exposed phosphate backbone of the ssDNA on the SWCNT. Moreover, higher content of negatively charged acidic amino acids led to enhancement in the protein corona (0.31 k_b_T / %acidic). This result indicates that protein charges are screened by salt in solution or that proteins interact directly with the solvent-accessible SWCNT surface. The latter hypothesis is supported by the low initial ssDNA coverage on the SWCNT and the small fraction of ssDNA removed from the SWCNT during protein adsorption.^34^ In addition, the latter hypothesis could explain the enhancement of aromatic residues, as they would then favorably interact via π-π stacking directly with the graphitic SWCNT surface.

(GT)_15_-SWCNTs have strikingly different interactions with proteins in CSF compared to plasma. In CSF, unlike in plasma, massive proteins are associated with favorable interactions with the SWCNTs (2.5 k_b_T / kDa) whereas more glycosylated proteins are associated with unfavorable interactions (−0.38 k_b_T / glycosylated site). Further in contrast with plasma, acidic residues in CSF are associated with unfavorable interactions with the nanoparticle (−0.52 k_b_T / %acidic), whereas positively charged arginine (and to a lesser extent, lysine) have favorable interactions (1.53 k_b_T / %R). The tendency of (GT)_15_-SWCNTs to interact favorably with positively charged residues and unfavorably with negatively charged residues suggests that the negatively charged (GT)_15_ ssDNA wrapping is less screened in CSF than in plasma.

Protein properties that are controlled for but that do not show a statistically significant effect on fold change for any nanoparticle in any biofluid include: the number of disulfide bonds (used as a proxy for protein stability), number of biomolecular binding sites, number of metal binding sites, and percentage of histidine or tryptophan. The lack of dependence on disulfide bond content and also instability index (see **Linear regression models** in Materials and Methods) is surprising in the context of previous corona literature, which suggests that less structurally stable proteins are more surface active.^50^

### Protein corona driving forces of formation

To gain further insight on interactions driving protein adsorption to nanoparticles, incubation conditions of (GT)_15_-SWCNTs exposed to plasma were varied and corona proteins were characterized by 2D PAGE. Specifically, conditions varied include dynamics (to probe corona stability), ionic strength (to probe electrostatic interactions), and temperature (to probe entropic contributions) (Figure 6). Under dynamic protein-nanoparticle incubation conditions, proteins in the outer adsorbed plane undergo shear and are removed. The proteins that remain are postulated to represent the “hard”, inner corona proteins that interact more strongly with the nanoparticle surface, whereby removed proteins represent the “soft”, outer corona proteins that interact with other adsorbed proteins. For the case of (GT)_15_-SWCNTs, apolipoproteins A-I, clusterin, complement C3, fibrinogen, and alpha-1-antitrypsin compose the hard corona, while some soft corona proteins of interest include albumin and haptoglobin. Elimination of salt (changing the buffer from 0.1 M PBS to water) increases the role of electrostatic interactions by removing ionic charge screening. More specifically, repulsive electrostatic forces originate from interactions between electric double layers surrounding the charged colloidal surfaces and proteins, which scale inversely with the square-root of salt ionic strength in solution. When incubated in water, the absence of charge screening means that proteins and nanoparticle surfaces do not approach as closely in solution, while lateral electrostatic interactions of surface-adsorbed proteins increase, both of which result in less protein adsorption.^51^ For (GT)_15_-SWCNTs, the three hard corona proteins of apolipoprotein A-I, complement C3, and fibrinogen still enter the corona despite the lack of charge shielding. This result points to the role of non-electrostatic forces facilitating formation of the hard corona, likely hydrophobic interactions, that drive protein-SWCNT adsorption even under electrostatically adverse conditions of electric double-layer repulsion (as supported by zeta potential measurements, **Figure S2**). In addition, most soft corona proteins, as determined by dynamic incubation conditions, are missing in the no salt incubation, reaffirming the need for charge-screening in the formation of the soft corona. In the final column of **Figure 6b**, higher temperature incubation increases weighting of the entropic contribution to the overall free energy change of binding. Entropic contributions originate from the solvent (positive, as the hydration shells of the protein and surface are released to bulk) and protein (negative, from the adsorbate losing degrees of freedom and potentially positive, as the protein may unfold upon adsorption). At high temperature, apolipoprotein A-I, complement C3, and fibrinogen are still able to adsorb to (GT)_15_-SWCNTs, indicating that adsorption of these proteins is entropically favorable and/or enthalpically driven. The interplay of binding entropy and enthalpy is investigated further in the corona dynamics section via application of isothermal titration calorimetry (ITC). Thus, apolipoprotein A-I, complement C3, and fibrinogen undergo high affinity binding to (GT)_15_-SWCNTs despite dynamic perturbation, low ionic strength, and increased temperature incubation conditions. We hypothesize that these proteins constitute the inner, hard corona around (GT)_15_-SWCNTs, with hydrophobic interactions driving the formation of the hard corona. The predominance of hydrophobic interactions in forming the hard corona suggests protein denaturation upon adsorption to SWCNTs, as most hydrophobic portions of proteins are buried and not intrinsically solvent-accessible.^52^ Our results suggest increasing SWCNT surface charge and employing SWCNT-based technologies in dynamic flow conditions (such as in circulating environments) will reduce outer-corona biofouling, whereas the inner corona will form despite high-shear flow conditions, although it will be reduced by increasing SWCNT surface hydrophilicity.

**Figure 6.**
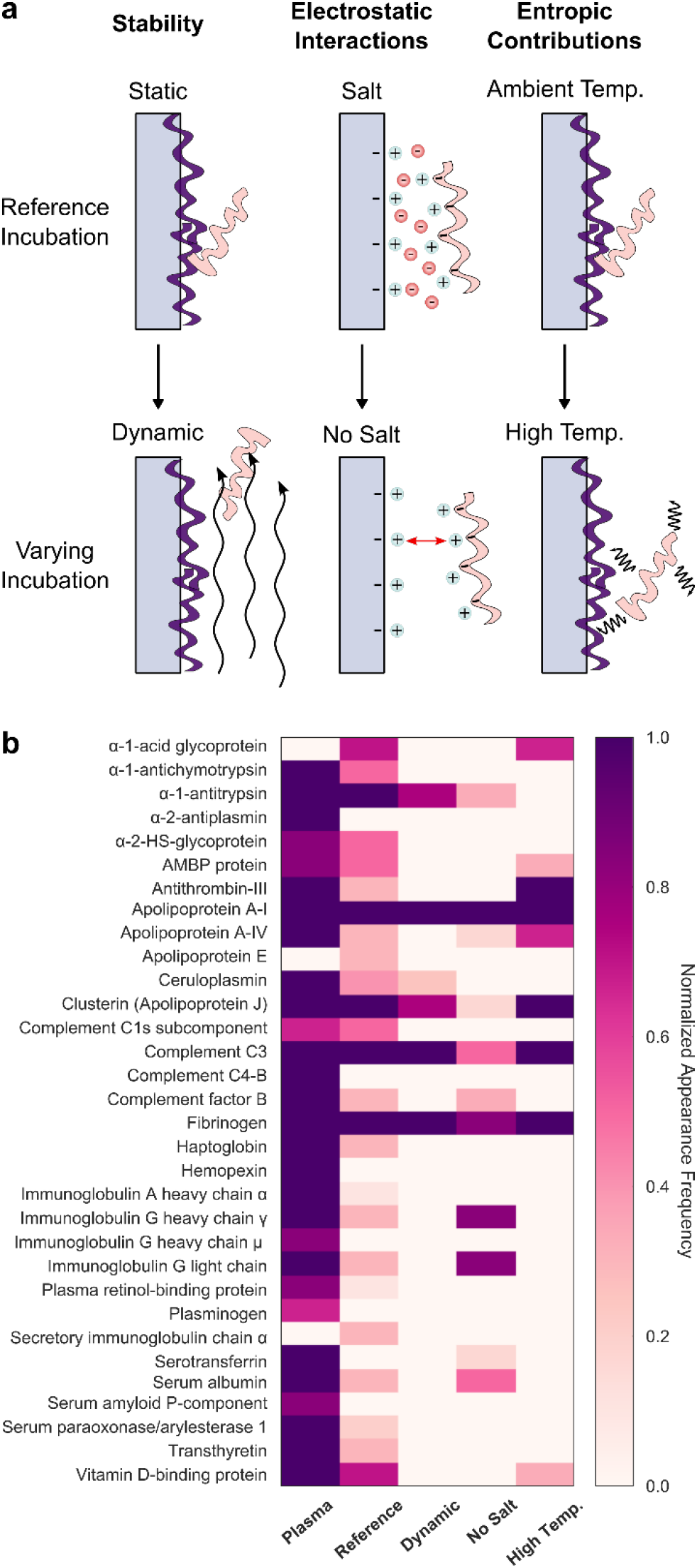
Effect of varying incubation parameters to probe corona stability, electrostatic interactions, and entropic contributions to corona formation of plasma proteins on (GT)_15_-SWCNTs. **(a)** Schematics depicting incubation conditions affecting corona adsorption, with reference conditions (top) vs. varied conditions (bottom). **(b)** Plasma proteins adsorbed on (GT)_15_-SWCNTs are compared to proteins present in the native plasma biofluid (left-most column) and the plasma corona formed on (GT)_15_-SWCNTs under reference conditions (static, 0.1 M phosphate-buffered saline, and 25°C incubation). Varying incubation conditions include: dynamic (on orbital shaker; to probe corona stability), no salt (water; to probe ionic effects), and high temperature (37°C; to probe entropic contributions). Color indicates normalized appearance frequency of protein in corona, as characterized by 2D PAGE.

### Protein corona dynamics

Proteins adsorb rapidly to nanoparticle surfaces, reportedly within 30 seconds of biofluid exposure.^15,43^ Beyond probing corona composition at the end-point of adsorption, we next investigated corona formation dynamics to understand the time-dependent process and overall system energetics driving corona formation. Thermodynamic and kinetic modes of probing protein adsorption to SWCNT surfaces were employed towards this end. Isothermal titration calorimetry (ITC) was applied to deconvolute the enthalpic vs. entropic contributions to the thermodynamics of protein-nanoparticle binding.^5,23,53^ ITC records time-resolved heats of binding, thus the binding enthalpy and a binding curve. These measurements allow fitting to extract the equilibrium constant and binding stoichiometry from the time-dependent binding curve, then calculation of the change in Gibbs free energy, and finally calculation of the change in entropy.

ITC was applied to study the binding of (GT)_15_-SWCNTs with two proteins identified by LC-MS/MS with opposite binding affinities: albumin was selected as a model low-binding protein and fibrinogen as a model high-binding protein. For each ITC run, 10 μL of 1.2 g/L protein was added for each of 24 injections from the syringe into 1 mL of 0.1 g/L (GT)_15_-SWCNTs in the cell (see further details in SI). ITC results are presented in **Figure S9**, again confirming that fibrinogen does adsorb to (GT)_15_-SWCNTs and albumin does not, as evidenced by the binding curve in the former and absence of changing heats upon injection in the latter. From ITC binding curves of fibrinogen with (GT)_15_-SWCNTs, the change in enthalpy is -565.2 kJ/mol and the change in entropy is -1.756 kJ/K-mol. These energies reveal that (i) fibrinogen binding is exothermic, and (ii) fibrinogen binding is overall entropically unfavorable, i.e., there is net ordering of the system. Reiterating the earlier discussion on entropy of binding, this negative change in entropy implies that the protein “ordering” contribution upon moving from free in solution to constrained on a surface outweighs the solvent “disordering” contribution upon desolvation. Enthalpic contributions arise from (i) noncovalent interactions between protein and the SWCNT surface and (ii) hydrogen bond formation within the bulk solvent as protein leaves solution to enter the adsorbed state. This favorable enthalpic term outweighs the net unfavorable entropic terms to ultimately drive formation as a spontaneous, energetically favorable process: the net change in free energy is -41.91 kJ/mol. However, these ITC results must be interpreted with the consideration that the equilibrium requirement for this thermodynamic analysis is not rigorously held (as discussed in SI).^2,33,43^ This binding profile shape for protein-surface adsorption processes often emerges as a result of adsorption-induced protein spreading/denaturation, reorientation, and aggregation as a function of bulk protein concentration, in contrast to originating from the dynamic equilibrium between the fluid and surface-adsorbed phases required for Langmuirian adsorption.^54–56^ Thus, although these binding curves confirm compositional findings of the relative binding affinities, it should be noted that ITC is not a suitable methodology to study all nanoparticle-protein systems and these limitations must be reflected in interpreting these binding energetics as overall system energies, rather than a true deconvolution of protein-nanoparticle binding interactions.

To further investigate the dynamics of protein corona formation, we implemented a real-time kinetic binding assay to study protein interactions with SWCNTs.^34^ Briefly, multiplexed fluorescence enables tracking each entity involved in the corona formation and exchange process, with cyanine 5 (Cy5)-tagged ssDNA originally on the SWCNT surface exchanging with protein added to solution. Our previous work confirms that fibrinogen adsorbs and displaces more ssDNA from the SWCNT surface than albumin, which only binds to a low degree and displaces a lesser amount of ssDNA. We implemented this platform to track the binding of key plasma corona proteins to (GT)_15_-SWCNTs and (GT)_6_-SWCNTs (**Figure 7a** and **Figure 7b**, respectively), with desorption of Cy5-tagged ssDNA originally on the SWCNT used as a proxy for protein adsorption to SWCNT. Specifically, we assayed the protein panel: clusterin, apolipoprotein A-I, fibrinogen, and complement C3, which are predicted to adsorb in high abundance to (GT)_15_-SWCNTs, and alpha-2-HS glycoprotein, immunoglobulin G, and albumin, which are predicted to adsorb less to (GT)_15_-SWCNTs based on LC-MS/MS compositional analysis (**Figure S4**). Based on the compositional data, we expect the ordering of molar protein amounts in the plasma corona to be: clusterin > apolipoprotein A-I > complement C3 > fibrinogen > immunoglobulin G > alpha-2-HS glycoprotein > albumin. Interestingly, the order of protein adsorption was: fibrinogen > apolipoprotein A-I > alpha-2-HS glycoprotein > immunoglobulin G ≈ clusterin > complement C3 > albumin > phosphate-buffered saline (PBS) control (**Figure 7a**). While this result affirms the high binding affinity of fibrinogen and apolipoprotein A-I vs. low binding affinity of albumin to (GT)_6_-SWCNTs, some of the single-protein end-states do not match the relative ordering of protein abundances from the full-biofluid LC-MS/MS experiments. This result further supports that higher order interactions, such as the Vroman effect, affect protein adsorption in the full-biofluid experiments, absent in the single-protein experiments. Moreover, these time-dependent dynamics reveal that the rates of protein binding are distinct among proteins, even though some converge to the same final value (such as alpha-2-HS glycoprotein vs. clusterin in **Figure 7a**). Note that in the case of the control, ssDNA adsorption to the SWCNT is observed, in line with previous studies.^34^ A comparison of this same protein panel binding to (GT)_6_-SWCNTs is provided because the shorter ssDNA strand is displaced more readily, providing a greater spread between protein species. For (GT)_6_-SWCNTs, the expected ordering of molar protein amounts in the plasma corona is: apolipoprotein A-I > complement C3 > clusterin > fibrinogen > immunoglobulin G > alpha-2-HS glycoprotein > albumin (**Figure S4**). The dynamics of protein adsorption recapitulate similar high- vs. low-binding propensity proteins, yet, again complement C3 and clusterin display significantly less adsorption than expected based on LC-MS/MS results, signifying that these particular proteins enter the corona with a cooperative binding mechanism, rather than simply displaying high binding affinity to the SWCNT surface on their own. Our dynamic tracking results show, in sum, that protein corona formation is a function of collective interactions at the nano-bio interface, rather than a property of isolated protein-nanomaterial surface interactions.^4^ To more quantitatively build a physical picture of protein-SWCNT association, we next expand to structural studies of these complexes.

**Figure 7.**
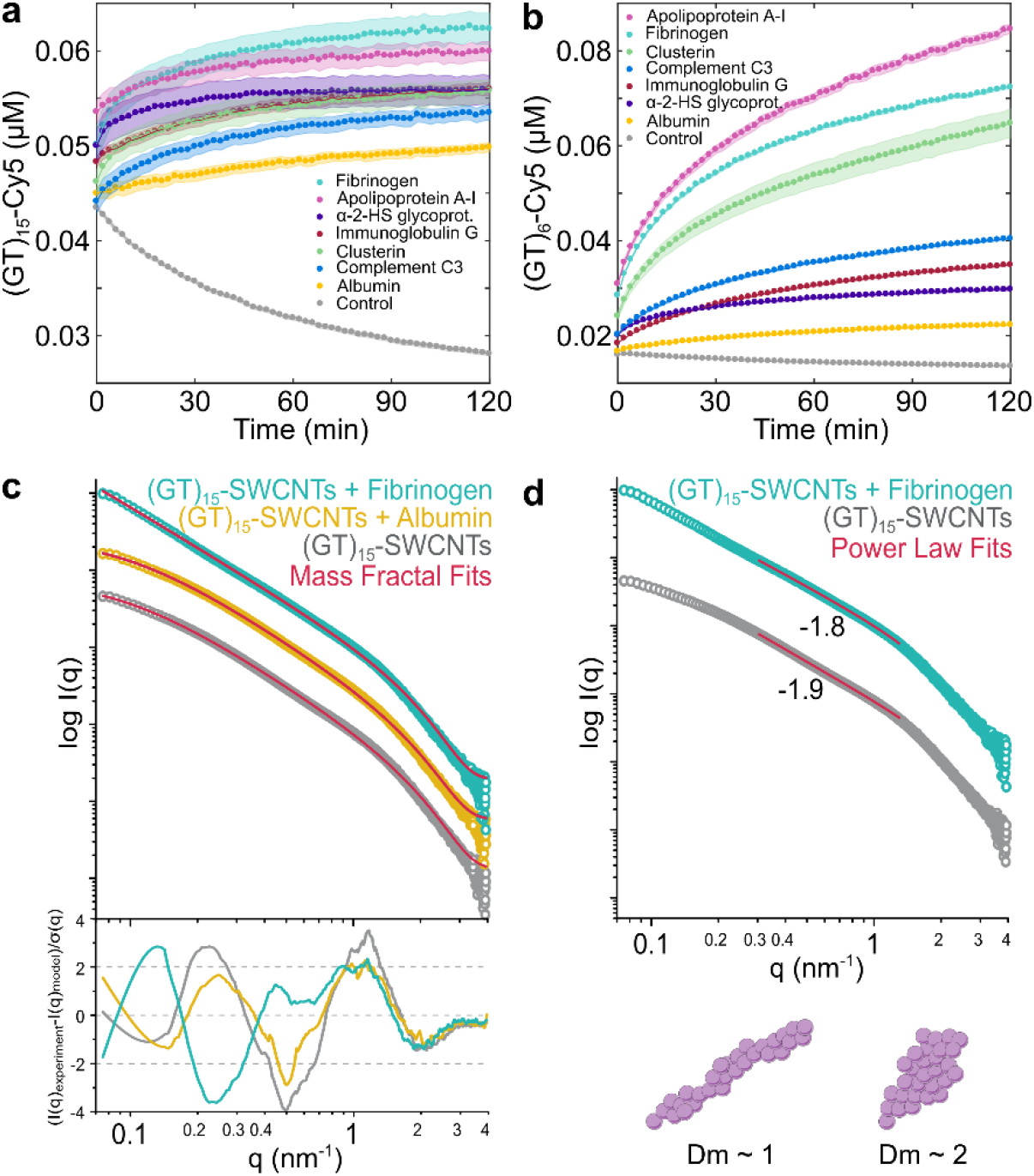
Protein corona dynamics and structure assessed for binding of key proteins to ssDNA-SWCNTs. A corona exchange assay is employed to determine binding kinetics of a protein panel (each at 80 mg/L final concentration) to **(a)** (GT)_15_-SWCNTs and **(b)** (GT)_6_-SWCNTs (5 mg/L final concentration). Small-angle x-ray scattering (SAXS) is applied to gain in-solution structural information of albumin vs. fibrinogen adsorption on (GT)_15_-SWCNTs. **(c)** Experimental small-angle x-ray scattering (SAXS) profiles for 0.5 g/L (GT)_15_-SWCNTs with and without albumin or fibrinogen, each at 0.5 g/L final concentrations. Mass fractal model fits are included in red together with fit residuals below. **(d)** SAXS profiles fit to show power law dependencies in the Porod regions, with illustration depicting the mass fractal dimension *Dm* increasing from approximately 1 to 2.

### Protein corona structure with small angle X-ray scattering

To evaluate structural changes in the (GT)_15_-SWCNTs due to protein corona formation, small angle X-ray scattering (SAXS) was performed with the two proteins, albumin and fibrinogen, as model low-binding and high-binding proteins, respectively. SAXS enables in-solution structural measurements, therefore we can assess the morphology of the protein-nanoparticle complex in an aqueous, more biologically relevant context. Solution SAXS probed the structure of 0.5 g/L (GT)_15_-SWCNTs in the presence of 0.5 g/L albumin or fibrinogen (**Figure 7** and **Figure S10**). The linear combination (GT)_15_-SWCNTs and albumin standard curves produced a SAXS profile identical to the mixed sample of (GT)_15_-SWCNTs+albumin, suggesting no interaction between the species. Dissimilarly, no calculated linear combination of the (GT)_15_-SWCNTs and fibrinogen standard curves could be produced to fit the SAXS profiles of the mixed sample, (GT)_15_-SWCNTs+fibrinogen, indicating formation of unique form factors and thus complexation. These mixing results recapitulate the corona composition results, where fibrinogen does adsorb to (GT)_15_-SWCNTs whereas albumin does not. Additionally, a clear concentration dependence is observed with an increase in the ratio of fibrinogen to (GT)_15_-SWCNTs by two-fold, while albumin shows no additional binding at elevated concentrations (**Figure S10a, Figure S10b**). Surface-adsorbed fibrinogen on (GT)_15_-SWCNTs was also confirmed via visualization with transmission electron microscopy (TEM; **Figure S11**).

All (GT)_15_-SWCNT samples with and without proteins were determined to be intrinsically disordered and experimental SAXS profiles were accordingly fit using mass fractal geometries (**Figure 7c**; see full derivation in SI). These models were complemented by calculating power-law dependencies from the Porod region, as demonstrated in **Figure 7d**. Three main values are derived from these mass fractal and power-law calculations: (i) radius *R* (nm), (ii) fractal dimension *Dm*, and (iii) cutoff length *ζ* (nm) (included in **Table S4**).^57–59^ The radius *R* in the mass fractal analysis is traditionally defined as the radius of the uniform sphere used to cover the fractal. These radii were approximately 1 nm for the analysis of all samples, suggesting that the overall topology of the (GT)_15_-SWCNTs remain constant with and without protein. The fractal dimension *Dm* and analogous power-law exponent *p* estimate the overall bulk geometries of the mass fractals, where the integer values of these variables represent the three dimensions in Euclidean space. Thus, *Dm* or *p* = 1, 2, or 3 represent rod, disk, or sphere geometries, respectively. The *Dm* and *p* values for (GT)_15_-SWCNTs and (GT)_15_-SWCNTs+albumin are |1.89, while (GT)_15_-SWCNTs+fibrinogen are ~ 1.77. These values suggest an initial disk-like geometry with slight rod-like character before binding to protein (or in the presence of albumin, which does not significantly bind), then the decrease in *Dm* in the case of (GT)_15_-SWCNT+fibrinogen suggests an elongation of the overall mass fractal structure (**Figure 7d**). This elongation is consistent with previous literature in which fibrinogen binds to SWCNTs in a lengthwise manner according to atomic force microscopy and molecular dynamics simulation,^60^ specifically with the two outer D-domains driving adsorption.^61^ Moreover, the decrease of (GT)_15_-SWCNT *Dm* in the presence of fibrinogen signifies increasing attractive forces between the molecular entities and according colloidal instability.^62^ Finally, the cutoff length *ζ* defines the maximum distance between any two points of the mass fractal. The calculated cutoff lengths show the most drastic increase from (GT)_15_-SWCNTs with and without albumin, ~11 nm, to (GT)_15_-SWCNTs with fibrinogen, ~100 nm, revealing a significant increase in the overall size of the aggregation. Thus, SAXS confirms fibrinogen complexation with (GT)_15_-SWCNTs, suggests a side-on orientation (as reiterated by TEM), and enables quantification of the changing fractal structure, pointing to the role of multilayer adsorption mechanisms and aggregate formation. More broadly, SAXS presents a valuable methodology to measure protein-nanoparticle complexation in solution, where knowledge of aggregate size and morphology is valuable in assessing nanoparticle interactions *in vivo*.

## Conclusions

Formation of the protein corona on engineered nanoparticles can lead to profound downstream effects within biological systems.^2,3^ Protein adsorption to the nanoparticle not only impacts the intrinsic structure and function of the proteins, but these adsorbed proteins also endow new biochemical properties to the nanoparticle and alter intended nanoparticle function.^2,6–10^ This phenomenon frequently results in undesirable nanoparticle outcomes including off-target localization, reduced efficacy, and toxicity.^2,9,11,12^ There is need to develop a more rigorous understanding of protein corona composition, driving forces of formation, dynamics, and structure, particularly on understudied nanoparticles such as SWCNTs and in biofluids such as cerebrospinal fluid. While prior studies clarify different aspects of bio-corona formation, system constraints such as surface-immobilization or treating the protein corona as existing at thermodynamic equilibrium make it difficult to reliably translate results to real biofluid systems.^2,33,43^ Accordingly, we have applied multimodal techniques to characterize the adsorbed protein corona in a more biologically representative, in-solution state.

In this work, we have developed and validated a platform to determine protein corona composition on two distinct nanoparticles, PNPs and (GT)_15_-SWCNTs, in two biofluids, blood plasma and cerebrospinal fluid. The framework itself is generic to study protein corona composition on other nanoparticles in other biofluids. We find that previously studied PNPs display a more agnostic character towards plasma proteins of various functional classes entering the corona, whereby the protein corona abundance scales with abundance within the bulk biofluid. Conversely, we find that high-aspect ratio (GT)_15_-SWCNTs instead show strong preferential binding of certain protein classes, with corona enrichment of plasma proteins involved in lipid transport, complement activation, and blood coagulation, and corona enrichment of CSF proteins involved in cell adhesion/signal transduction, blood coagulation, and complement activation. The selectivity of proteins binding to (GT)_15_-SWCNTs is a promising result in motivating the development of SWCNT-based nanosensors passivated against uncontrolled biofouling within protein-rich environments. Of note, (GT)_15_-SWCNTs show high-abundance and enriched binding of fibrinogen, and low-abundance and depleted binding of albumin. This raises a cogent concern for the status quo of testing nanotechnologies in blood serum (absent of fibrinogen) rather than blood plasma (with all protein constituents present), where fibrinogen may be a more important contributor to diminished *in vivo* efficacy than albumin.

With this consensus of corona composition, protein molecular attributes that dictate protein-nanoparticle interactions are interrogated by regression analysis of the ln-fold change in corona composition as compared to the native biofluid. These regression coefficients present a connection between the protein properties and the thermodynamic free energy change of proteins binding to the nanoparticle. The presence of aromatic amino acids largely leads to enhanced interactions of both plasma and CSF proteins with (GT)_15_-SWCNTs, highlighting the contribution of π-π interactions of protein adsorbates with the graphitic SWCNT surface. Characteristics of plasma proteins adsorbed to (GT)_15_-SWCNTs demonstrate deviations from CSF proteins: higher positively charged residue content is associated with unfavorable interactions and higher negatively charged residue content with favorable interactions in plasma, while the opposite is true in CSF. Corona composition formed under variable incubation conditions indicates that hydrophobic interactions play a key role in formation of the hard corona on (GT)_15_-SWCNTs (proteins interacting more directly with the nanoparticle surface with longer corona residence times), while electrostatic screening is implicated in formation of the shear-sensitive soft corona (proteins predominantly interacting with other adsorbed proteins that undergo more exchange with surrounding bulk-phase proteins).

Study of transient corona thermodynamics and kinetics further supports differential protein affinity for ssDNA-SWCNTs and underscores the complexity of corona formation via higher-order interaction mechanisms including the Vroman effect and post-adsorptive transitions. These results are significant in emphasizing the non-intuitive nature of corona formation, in that seemingly important properties for corona formation such as protein stability or specific binding sites are not predictive of presence in a nanoparticle corona.

This work clarifies fundamental interactions that can be applied to nanoscale systems in which the development and optimization is done *in vitro*, but with a desired end application *in vivo*. A more in-depth understanding of the protein corona could allow *a priori* prediction of biodistribution profiles and/or enable us to better understand these results in organisms. Moreover, outcomes of this work will inform rational nanoparticle design to either mitigate corona formation and preserve the as-designed nanoparticle, or to promote adhesion of specific biomolecules to favor accumulation in targeted areas within the whole organism. Elucidating protein corona composition, dynamics, structure, and driving forces that mediate nanoparticle-protein interactions will establish design considerations for nanosensor development and provide a framework for understanding *how* and *why* our engineered nanoparticles are affecting, and being affected by, complex bioenvironments.

## Materials and Methods

### Synthesis of SWCNT-based nanosensors

Single-stranded DNA with single-walled carbon nanotube (ssDNA-SWCNT) suspensions were prepared with 1 mg of mixed-chirality SWCNTs (small diameter HiPco™ SWCNTs, NanoIntegris) and 1 mg of ssDNA (custom ssDNA oligos with standard desalting, Integrated DNA Technologies, Inc.) in 1 mL of 0.1 M phosphate-buffered saline (PBS). Solutions were bath sonicated for 10 min (Branson Ultrasonic 1800) and probe-tip sonicated for 10 min in an ice bath (3 mm probe tip at 50% amplitude, 5-6 W, Cole-Parmer Ultrasonic Processor). Samples were incubated at room temperature for 30 min then centrifuged to pellet insoluble bundles and contaminants (16.1 krcf for 30 min). Supernatant containing the product was collected. ssDNA-SWCNTs were spin-filtered to remove free ssDNA (Amicon Ultra-0.5 mL centrifugal filters with 100 kDa MWCO, Millipore Sigma) by washing with Milli-Q water two times (8 krcf for 5 min) then reversing the spin filter and centrifuging to recover sample (1 krcf for 5 min). ssDNA-SWCNT concentration was determined via sample absorbance at 632 nm (NanoVue Plus, GE Healthcare Life Sciences) and the extinction coefficient ε_632nm_=0.036 L cm mg^−1^.^25^ ssDNA-SWCNTs were stored at 4°C until use and then diluted to a working concentration of 100 mg/L in 0.1 M PBS.

### Modified pull-down assay

Protein corona composition was studied by a modified pull-down assay (**Figure 1**) with three nanoparticles of interest: 100 nm diameter polystyrene nanoparticles (PNPs; Fluoresbrite® yellow-green fluorophore-labeled with ex441nm em486nm, Polysciences Inc.), (GT)_15_-SWCNTs, and (GT)_6_-SWCNTs. PNPs were vortexed prior to use (1 min in 5 s pulses) to ensure homogeneously dispersed solution. Biofluids studied were human plasma and human cerebrospinal fluid (CSF) (see details in **Table S5**). Biofluids were thawed once for aliquoting then stored at stored -80°C until use. CSF was concentrated 10X prior to the incubation step to maintain the same protein to nanoparticle ratios under volume constraints (14 krcf for 30 min; Amicon Ultra-0.5 mL centrifugal filters with 3 kDa MWCO, Millipore Sigma). The ratio of protein concentration to nanoparticle surface area was maintained constant for each respective nanoparticle type in different biofluids, with 26 g/L protein per m^2^ nanoparticle surface area for PNPs (based on previous literature^17^) and 200 g/L protein per m^2^ nanoparticle surface area for (GT)_6_-and (GT)_15_-SWCNTs (based on experimental optimization for both plasma and CSF). This translates to final concentrations of 1.67 g/L PNPs or 28.7 mg/L ssDNA-SWCNTs with 3.2% v/v plasma and 0.50 g/L PNPs or 10 mg/L ssDNA-SWCNTs with 6.4% v/v of 10X CSF. Nanoparticles were incubated with biofluid in 0.1 M PBS in 750 μL total volume for 1 h at ambient temperature. Protein-nanoparticle complexes were pelleted out of solution by centrifugation (16.1 krcf for 20 min). Supernatant containing unbound proteins was removed to waste, the pellet resuspended in 0.1 M PBS to the same volume, and the pellet was broken up by pipetting. Washing was repeated three times to ensure removal of unbound proteins.

### Two-dimensional polyacrylamide gel electrophoretic separation (2D PAGE)

2D PAGE was performed to identify proteins via separation by isoelectric point in the first dimension and molecular weight in the second dimension. For analysis by 2D PAGE, bound proteins were eluted from nanoparticles by heating at 95°C for 10 min in SDS/BME reducing buffer (2% SDS, 5% β-mercaptoethanol, 0.066 M Tris-HCl). 1D separation was run according to the O’Farrell protocol^63^ (adapted for Bio-Rad Mini-PROTEAN Tube Cell). Briefly, 1D sample buffer (8 M urea, 2% Triton X-100, 5% β-mercaptoethanol, 2% total carrier ampholytes -1.6% Bio-Lyte 5/7, 0.4% Bio-Lyte 3/10) was added to samples in a 1:1 or 0.07:1 volume ratio (relative to initial plasma and CSF volumes, respectively) and incubated for 10 min. 1D separation was carried out in capillary tube PAGE with gel composition of 4% acrylamide (total monomer), 8 M urea, 2% Triton X-100, 2% total carrier ampholytes, 0.02% ammonium persulfate (APS), and 0.15% Tetramethylethylenediamine (TEMED). 25 μL sample and 25 μL 1D sample overlay buffer (4 M urea, 1% total carrier ampholytes) was loaded per capillary tube gel. Upper and lower chamber buffers were 100 mM sodium hydroxide and 10 mM phosphoric acid, respectively. 1D separation was run at 500 V for 10 min, 750 V for 3.5 h. Nanoparticles were filtered from the eluted proteins by the gel itself. Capillary gels were extruded and loaded onto 2D gels. 2D separation was run according to the Laemmli protocol^64^ (adapted for Bio-Rad Mini-PROTEAN Tetra Cell). Briefly, SDS/BME reducing buffer was added to the 2D well to cover the capillary gel and incubated for 10 min. 2D separation was carried out in 1 mm vertical mini gel format with a discontinuous buffer system under denaturing conditions. Gel composition was 12% acrylamide (total monomer), 0.375 M Tris-HCl, 0.1% SDS, 0.05% APS, 0.05% TEMED for the resolving gel and 12% acrylamide (total monomer), 0.125 M Tris-HCl, 0.1% SDS, 0.05% APS, 0.1% TEMED for the stacking gel. Electrode buffer was 25 mM Tris, 192 mM glycine, and 3.5 mM SDS (pH 8.3). 2D separation was run at 200 V for 1 h. Gels were extracted and silver stained according to Bio-Rad’s Silver Stain Plus protocol and identified with ExPASy’s SWISS-2DPAGE database.^65^

### Liquid chromatography-tandem mass spectrometry (LC-MS/MS)

For analysis by LC-MS/MS, the modified pull-down protocol was performed with a scale-up by factors of 2, 4, 3, and 8 for PNPs/plasma, (GT)_x_-SWCNTs/plasma, PNPs/CSF, and (GT)_15_-SWCNTs/CSF, respectively, to ensure enough protein mass for analysis. Samples were prepared in experimental triplicate for plasma samples or run in technical triplicate for CSF samples. All spin filters were pre-rinsed according to manufacturer’s instructions to prevent sample contamination.

Bound proteins were eluted from nanoparticles by heating at 37°C for 60 min in urea/DTT reducing buffer (8 M urea, 5 mM DTT, 50 mM Tris-HCl, pH 8). Elution buffer was modified from SDS/βME for 2D PAGE to urea/DTT for LC-MS/MS analysis due to SDS interference with trypsin digestion, reverse-phase HPLC purification, and electrospray ionization efficiency.^66^ The profile of eluted proteins was confirmed to be invariable to the elution system by 2D PAGE, however, the total amount of eluted protein decreases. After elution from nanoparticles, protein concentration was determined with the EZQ Protein Quantitation Kit (Thermo Fisher Scientific). Eluted protein solution was centrifuged to remove nanoparticles (16 krcf for 20 min) and spin-filtered to concentrate and remove impurities (14 krcf for 30 min; Amicon Ultra-0.5 mL centrifugal filters with 3 kDa MWCO, Millipore Sigma). Proteins were alkylated with 15 mM iodoacetamide for 30 min in the dark. DTT was added to quench excess iodoacetamide in a volume ratio of 3:1 and incubated for 20 min. The reaction was diluted 1:1 with 50 mM Tris-HCl pH 8 to allow for enzymatic protein digestion. In-solution protein digestion was carried out in a ratio of 1:25 w/w Trypsin/Lys-C (Mass Spectrometry Grade, Promega) to protein overnight at 37°C. Nanoparticles were removed by spin filtering (14 krcf for 30 min; Amicon Ultra-0.5 mL centrifugal filters with 30 kDa MWCO, Millipore Sigma). Nanoparticle removal was done after protein digestion into peptides due to the otherwise very similar sizes of nanoparticles and proteins. Complete nanoparticle removal was verified with absorbance, fluorescence, and DLS measurements during method optimization. Peptide concentration was determined with the Pierce Peptide Quantitation Kit (Thermo Fisher Scientific) and all samples were normalized to 0.1 g/L in 100 μL total volume. Peptide solutions were spiked with 50 fmol of E. coli housekeeping peptide (Hi3 Ecoli Standard, Waters) per 5 μL sample volume to allow for protein quantification. Digestion was terminated by freezing samples to −20°C.

Proteolytically digested proteins were analyzed using a Synapt G2-S*i* mass spectrometer equipped with a nanoelectrospray ionization source and connected directly in line with an Acquity M-class ultra-performance liquid chromatography system (UPLC; Waters, Milford, MA). This instrumentation is in the California Institute for Quantitative Biosciences (QB3)/College of Chemistry Mass Spectrometry Facility at UC Berkeley. Data-independent, ion mobility-enabled mass spectra and tandem mass spectra^67–69^ were acquired in the positive ion mode. Data acquisition was controlled with MassLynx software (version 4.1) and tryptic peptide identification and quantification using a label-free approach^70–72^ were performed with Progenesis QI for Proteomics software (version 4.0, Waters).

Note that elution of proteins from nanoparticles with 5% SDS, 50mM TEAB in combination with the S-trap mini column purification methodology (Protifi; following manufacturer’s instructions) was attempted, however, this protocol was not pursued due to no improvement in protein sequence coverage and higher sample preparation times and material costs.

### Linear regression models

We linearly regressed^73^ the natural log of the fold change of proteins for each nanoparticle-biofluid pairing using two sets of protein descriptors. The first set of descriptors are categorical variables denoting what class a protein is in (*i.e.* 1 for a protein in a given class and 0 otherwise), namely, involved in acute-phase response, blood coagulation, cell adhesion/signal transduction, complement activation, immune response, lipid binding/transport, regulation of biological processes, transport, or miscellaneous/unknown (**Figure 3** and **Figure S7**; grouped according to PANTHER^39^). The variables were sum-effect coded such that the coefficients quantify how a protein class deviates from the grand mean of all protein classes and the intercept of the regression is the grand mean. Because each protein is grouped into one and only one class, the categorical variables are not linearly independent and one class is excluded from the regression;^73^ we chose the miscellaneous class.

The second set of descriptors are molecular and biophysical properties of the proteins: protein mass, fraction of amino acids that are non-aromatic hydrophobic (sum of alanine, valine, isoleucine, leucine, and methionine content), hydrophilic (sum of serine, threonine, asparagine, glutamine content), arginine (R), histidine (H), lysine (K), acidic (sum of aspartic acid and glutamic acid content), phenylalanine (F), tyrosine (Y), tryptophan (W), number of glycosylated sites, number of ligand binding sites, number of metal binding sites, and number of disulfide binds (**Figure 5** and **Figure S8**). Each of these descriptors is a continuous variable. The regression coefficients quantify the fractional difference in the fold change for a unit increase in the independent variable.^73^ Protein-specific information was acquired from UNIPROT.^48^ Note that these particular descriptors were chosen after primary analyses that eliminated highly co-dependent descriptors. An example was choosing to include percentage of acidic/basic amino acids rather than protein isoelectric point (from ExPASy Compute pI/MW), where the isoelectric point was deemed less exact because it relies on a theoretical calculation, omits protein fragments, and necessitates an average value for multicomponent proteins. Other examples were including number of disulfide bonds as an estimate of protein stability rather than protein instability index and segmenting to percentage of hydrophobic/aromatic amino acids rather than grand average hydropathy (GRAVY) score, in both cases due to the involvement of arbitrarily set scales (from ExPASy ProtParam).

For each regression, we included the measured protein fold changes for each replicate of a nanoparticle-biofluid system and controlled for sample-to-sample variability by including a categorical variable for the specific replicate. Protein abundances that fell below the lower limit of detection in the samples from the protein corona were set to 1×10^−5^ fmol, corresponding to the lowest detected protein abundance of all systems. Left-censoring the data in this way provides a conservative estimate of the regression coefficients by underestimating of the magnitude and significance.^73^ Calculated variance inflation factors for all variables in each independent regression was >4, indicating negligible multicollinearity between the independent variables.^74^ To avoid overestimating the statistical significance of independent variables, p-values were adjusted using the Benjamini-Hochberg false discovery rate procedure.^75^ All statistical analysis was implemented in Python using the StatsModels V0.10.1 package^76^ (0.27-0.39). **Table S2** and **Table S3** provide coefficients, standard errors, p-values, and R-squared values for each regression. The median R-squared of the first and second regression models for the nanoparticle-biofluid systems are 0.29 and 0.34, respectively, indicating the statistical models are descriptive rather than predictive. Nonlinear or decision tree algorithms provide more precise prediction of corona composition,^49^ however, these approaches were not considered because they are not readily interpretable, which is a principle goal of our analysis.

### Modified pull-down assay with varying incubation parameters

Protein corona composition was studied under varied incubation conditions in comparison to the reference state incubation (static, 0.1 M phosphate-buffered saline, and 25°C incubation). Varying incubation conditions include dynamic (on orbital shaker at medium speed; to test corona stability), no salt (water; to test ionic effects), and high temperature (37°C; to test entropic contributions). **Figure 6** summarizes data from 2D PAGE gels, with experimental replicates of 6 for plasma alone, 10 for reference, 4 for dynamic, 6 for no salt, and 3 for high temperature.

### Corona dynamics assay

Corona dynamic studies were completed as described previously.^34^ Briefly, the same suspension protocol was employed for preparation of fluorophore-labeled ssDNA-SWCNT complexes, using ssDNA-Cy5 (3’ Cy5-labeled custom ssDNA oligos with HPLC purification, Integrated DNA Technologies, Inc.) in place of unlabeled ssDNA. Lyophilized proteins were purchased (see details in **Table S5**) and reconstituted by adding 5 mg to 1 mL of 0.1 M PBS, tilting to dissolve for 15 min, filtering with 0.45 μm syringe filter (cellulose acetate membrane, VWR International), and quantifying with the Qubit Protein Assay (Thermo Fisher Scientific). Because of variation in amine-labeling of proteins, fluorescently labeled ssDNA was solely tracked, and the displacement of ssDNA from the SWCNT surface was taken as a proxy for protein adsorption. Equal volumes of 10 mg/L (GT)_15_- or (GT)_6_-Cy5-SWCNTs and 160 mg/L protein were added to a 96-well PCR plate (Bio-Rad) to a total volume of 50 μL. The plate was sealed with an optically transparent adhesive seal (Bio-Rad) and spun down on a benchtop centrifuge. Fluorescence time series readings were taken in a Bio-Rad CFX96 Real Time qPCR System, scanning the Cy5 channel every 2 min at 22.5°C. Fluorescence time series were analyzed without default background correction. Fluorescence values were converted to mass concentration using linear standard curves for ssDNA-Cy5.

### Small-angle X-ray scattering (SAXS)

SAXS data was collected at the SIBYLS beamline (bl12.3.1) at the Advanced Light Source of Lawrence Berkeley National Laboratory, Berkeley, California.^77^ X-ray wavelength was set at *λ* = 0.1127 nm and the sample-to-detector distance was 2.1 m, resulting in scattering vector (*q*) ranging from 0.1 -4 nm–1. The scattering vector is defined as *q* = 4πsin*θ*/*λ*, with scattering angle 2*θ*. Data was collected using a Dectris PILATUS3X 2M detector at 20°C and processed as described previously.^78^

Immediately prior to data collection, samples were added to 96-well plates kept at 10°C, where 30 μL of sample was bracketed with two wells of 30 μL 0.1 M PBS. Samples were transferred in the 96-well plate to the sampling position via a Tecan Evo liquid handling robot with modified pipetting needles acting as sample cells as described previously.^79^ Samples were exposed to X-ray synchrotron radiation for 30 s at a 0.5 s frame rate for a total of 60 images. For each sample collected, two 0.1 M PBS buffer samples were also collected to reduce error in subtraction. Each collected image was circularly integrated and normalized for beam intensity to generate a one-dimensional scattering profile by beamline-specific software. Buffer subtraction was performed for the one-dimensional scattering profile of each sample using each of the two buffer wells to ensure the subtraction process was not subject to instrument variations. Scattering profiles over the 30 s exposure were sequentially averaged together to eliminate any potential radiation damage effects. Averaging was performed with web-based software FrameSlice (sibyls.als.lbl.gov/ran). Mass fractal modeling details are included in SI.

## Acknowledgements

Proteomic mass spectrometry data collection at the California Institute for Quantitative Biosciences (QB3)/College of Chemistry Mass Spectrometry Facility for LC-MS/MS was funded through grant 1S10OD020062-01. SAXS data collection at the SIBYLS beamline (bl12.3.1) was funded through DOE BER Integrated Diffraction Analysis Technologies (IDAT) program and NIGMS grant P30 GM124169-01, ALS-ENABLE. This work benefited from the use of the SasView application, originally developed under NSF award DMR-0520547. SasView contains code developed with funding from the European Union’s Horizon 2020 research and innovation programme under the SINE2020 project, grant agreement No 654000. We thank Behzad Rad, Blakely Tresca, and Ron Zuckerman for assistance with ITC at the Molecular Foundry, supported by the Office of Science, Office of Basic Energy Sciences, of the U.S. Department of Energy under Contract No. DE-AC02-05CH11231. We thank Younghun Sung for TEM images acquired at the National Center for Electron Microscopy through proposal #5501, working with Dr. Karen Bustillo. We thank Matt Francis for Zetasizer Nano use. We acknowledge support of an NIH NIDA CEBRA award # R21DA044010 (to M.P.L.), a Burroughs Wellcome Fund Career Award at the Scientific Interface (CASI) (to M.P.L.), the Simons Foundation (to M.P.L.), a Stanley Fahn PDF Junior Faculty Grant with Award # PF-JFA-1760 (to M.P.L.), a Beckman Foundation Young Investigator Award (to M.P.L.), and a DARPA Young Investigator Award (to M.P.L.). M.P.L. is a Chan Zuckerberg Biohub investigator. R.L.P. and D.Y. acknowledge the support of NSF Graduate Research Fellowships (NSF DGE 1752814).

## Author Contributions

R.L.P. and M.P.L. conceived the idea and designed the study. R.L.P. performed the majority of the experiments and data analysis, and wrote the manuscript. D.Y. assisted in experimental design for and running of corona exchange assays. D.J.R. completed and analyzed SAXS results together with M.H. T.C. performed 2D PAGE under variable incubation conditions. A.R.C. modeled protein corona LC-MS/MS results. A.T.I. performed LC-MS/MS. All authors discussed the results, edited the manuscript, and approved the final version.

