## Supplemental Information for "Protein Corona Composition and Dynamics on Carbon Nanotubes in Blood Plasma and Cerebrospinal Fluid"

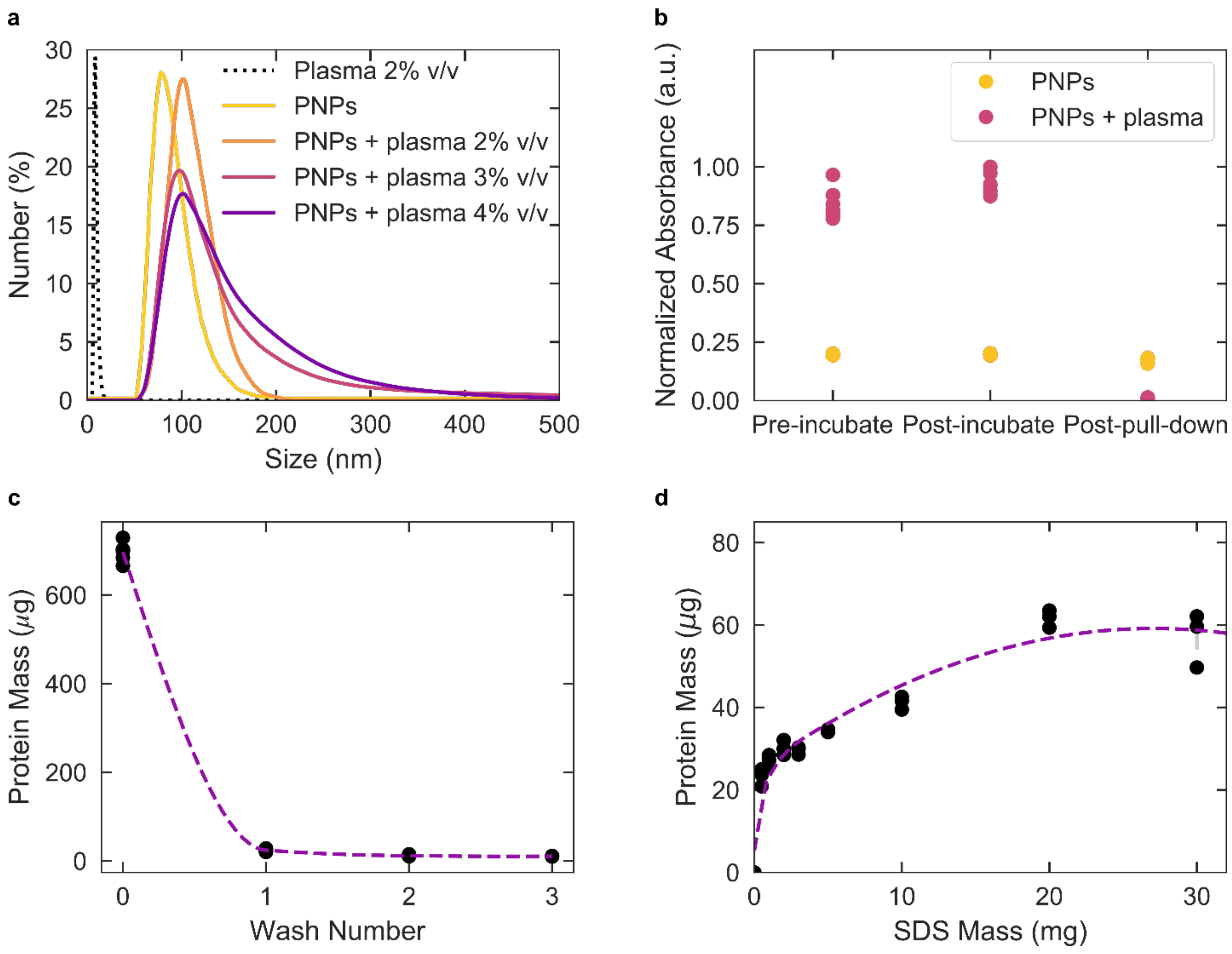


**Figure S1. Modified pull-down assay validation. (a)** Dynamic light scattering (DLS) reveals that plasma protein corona formation induces an increase in the hydrodynamic radius of the PNPs (1.25 g/L in 0.1 M PBS) via peak shifting and broadening. **(b)** Absorbance at PNP excitation max (441 nm) immediately after adding plasma to incubation solution ('initial'), incubating for 1 hour ('post-incubation'), and after the first pelleting step ('post-pull-down') demonstrates the presence of proteins facilitates pull-down of nanoparticles from solution in the initial pelleting step. **(c)** Quantification of free protein in solution via Qubit Protein Assay for varying wash number shows nearly complete depletion of free protein by three washes. **(d)** Quantification of eluted protein from nanoparticles via Pierce 660 nm Protein Assay with increasing SDS reducing buffer confirms complete elution of bound proteins from nanoparticle surface prior to characterization. Error bars on (b)-(d) are ± standard error for experimental replicates of N = 6, 6, and 3, respectively.

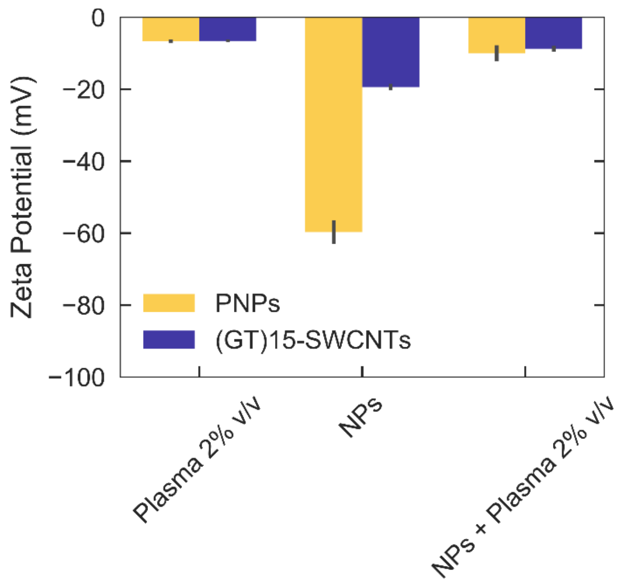


**Figure S2**. **Surface charge changes induced by plasma protein corona formation.** Zeta potential of native plasma, nanoparticles alone (PNPs yellow, $(GT)_{15}$-SWCNTs purple), and plasma protein-nanoparticle complexes. Lower magnitude zeta potential of protein-nanoparticle complexes indicates reduction in colloidal stability in the presence of surface-adsorbed proteins, as expected by visible aggregates formed.

#### Validation of modified pull-down assay

Each step was validated for polystyrene nanoparticles (PNPs) exposed to blood plasma proteins as follows: (i) incubation of proteins with nanoparticles induced an increase in nanoparticle hydrodynamic radius as determined by dynamic light scattering (DLS), where the number distribution both shifted to a larger peak center and broadened out due to nonuniform aggregate formation, as protein to nanoparticle loading was increased (**Figure S1**a); (ii) proteins initiated nanoparticle aggregation, thus facilitating nanoparticle pelleting and recovery for analysis, as shown by solution absorbance (Tyndall effect) before and after initial pull-down (**Figure S1**b); (iii) three washing steps were sufficient to remove unbound proteins by quantifying proteins remaining in the supernatant (**Figure S1**c); and (iv) proteins were fully eluted from nanoparticles by boiling in solutions of sodium dodecyl sulfate/β-mercaptoethanol (SDS/βME, for 2D PAGE analysis; **Figure S1**d) and urea/dithiothreitol (urea/DTT, for LC-MS/MS analysis). The equivalent verification of the pull-down assay was performed with $(GT)_{15}$-SWCNTs, yet the high aspect ratio of SWCNTs prohibited accurate DLS measurement.

Towards (i), protein adsorption to the nanoparticle surface during the incubation step was characterized by dynamic light scattering and zeta potential measurements in folded capillary zeta cell disposable cuvettes (Zetasizer Nano, Malvern Panalytical). Note that PNPs are negatively charged as a result of initiator fragments from the polymerization process, yet these PNPs are conventionally considered to be a model plain nanoparticle due to no explicit functionalization.[1] $(GT)_{15}$-SWCNTs are slightly negatively charged due to the presence of the ssDNA on the surface, with the phosphate backbone extending into solution. Towards (ii), absorbance spectra for pull-down verification were measured in a 700 µL volume, black-sided quartz cuvettes (Thorlabs, Inc.) with a UV-VIS-nIR spectrophotometer (Shimadzu UV-3600 Plus). For (iii), free protein remaining in the supernatant after pull-down was quantified during subsequent wash steps using the Qubit Protein Assay (Thermo Fisher Scientific). For (iv), eluted protein from the nanoparticle was quantified using the Pierce 660nm Assay (with Ionic Detergent Compatibility Reagent; Thermo Fisher Scientific). The denaturant/reducing agent combination SDS/BME was chosen for 2D PAGE by convention and urea/DTT due to the incompatibility of detergents with LC-MS/MS systems. While urea/DTT eluted less total protein mass in comparison to SDS/βME, there were no qualitative differences in protein composition based on both 2D PAGE and S-trap LC-MS/MS analysis.

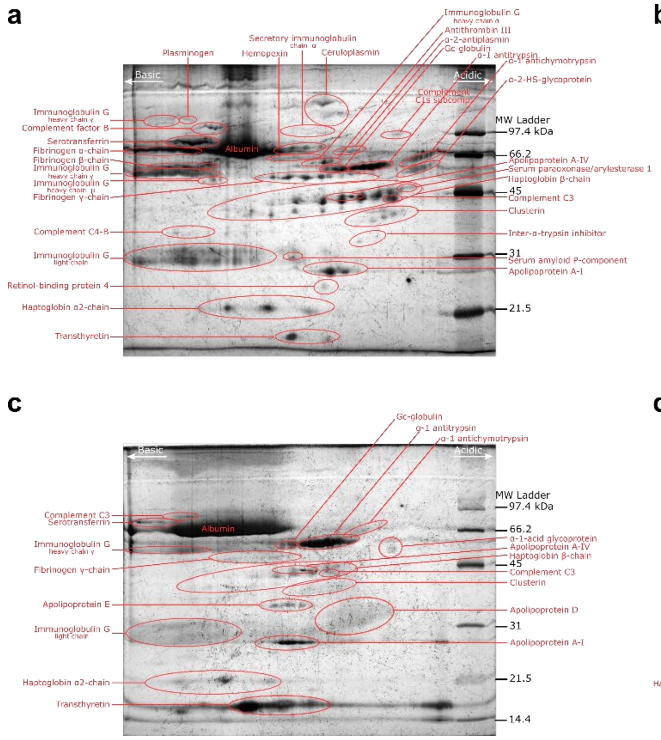


**Figure S3. Representative 2D PAGE gels. (a)** Plasma alone, **(b)** Plasma protein corona composition formed on $(GT)_{15}$-SWCNTs, **(c)** CSF alone, and **(d)** CSF protein corona composition formed on PNPs.

**a**

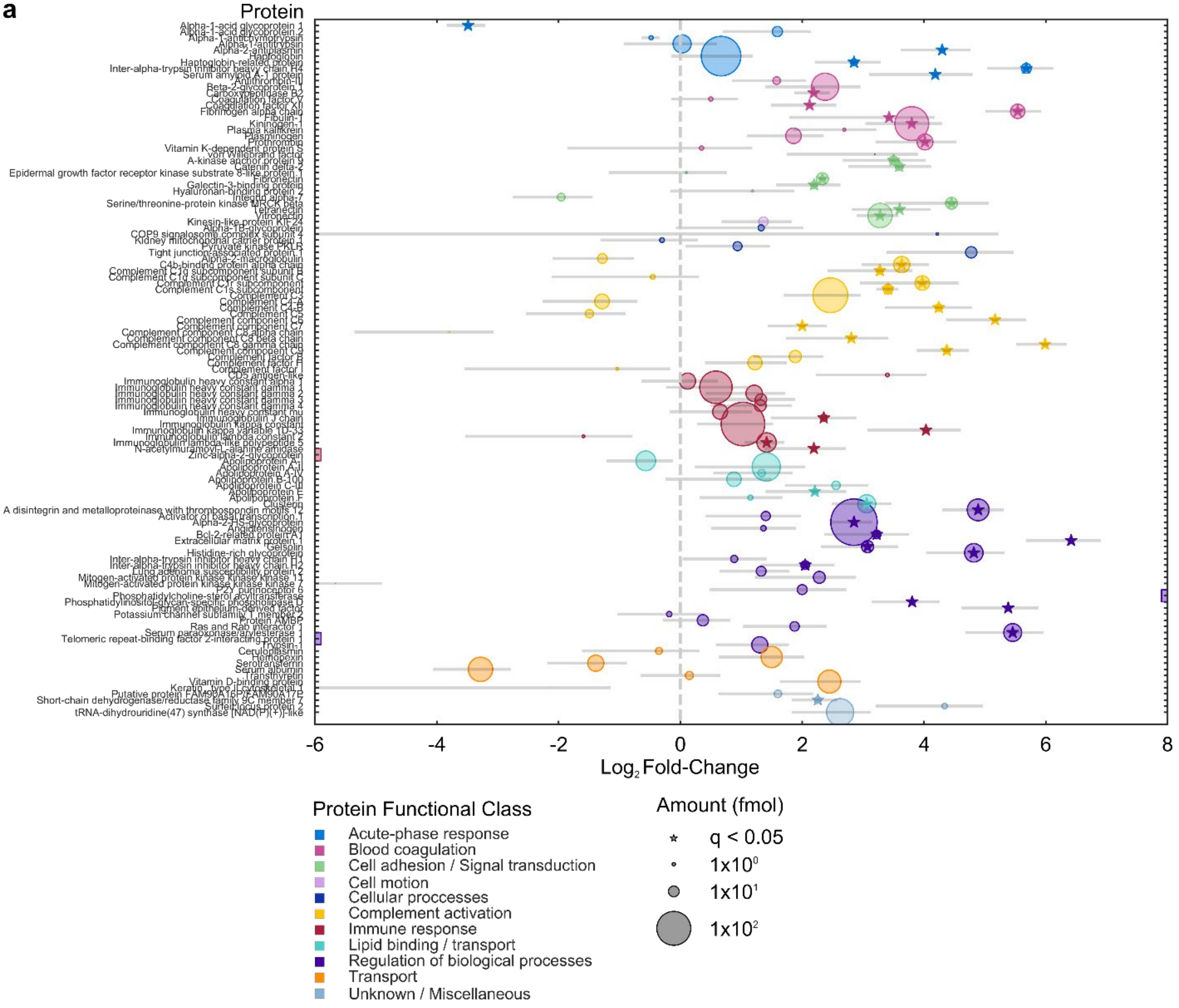
Protein
-6
-4
-2
0
2
4
6
8
Log2 Fold-Change
Protein Functional Class
Acute-phase response
Blood coagulation
Cell adhesion / Signal transduction
Cell motion
Cellular proccesses
Complement activation
Immune response
Lipid binding / transport
Regulation of biological processes
Transport
Unknown / Miscellaneous
Amount (fmol)
q < 0.05
1x10^0
1x10^1
1x10^2

**b**

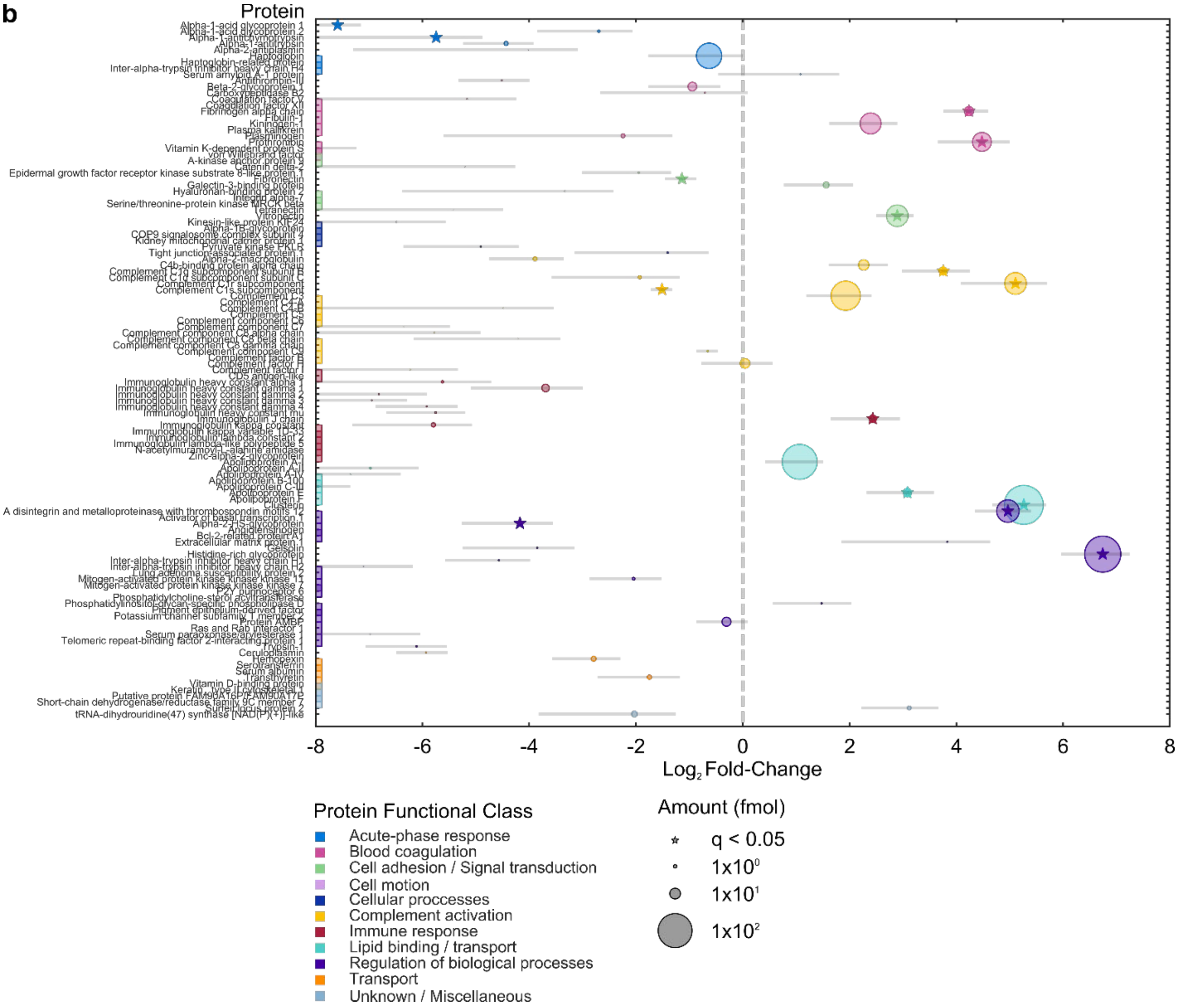
Protein
Log₂ Fold-Change
-8
-6
-4
-2
0
2
4
6
8
Protein Functional Class
Acute-phase response
Blood coagulation
Cell adhesion / Signal transduction
Cell motion
Cellular proccesses
Complement activation
Immune response
Lipid binding / transport
Regulation of biological processes
Transport
Unknown / Miscellaneous
Amount (fmol)
q < 0.05
1x10⁰
1x10¹
1x10²

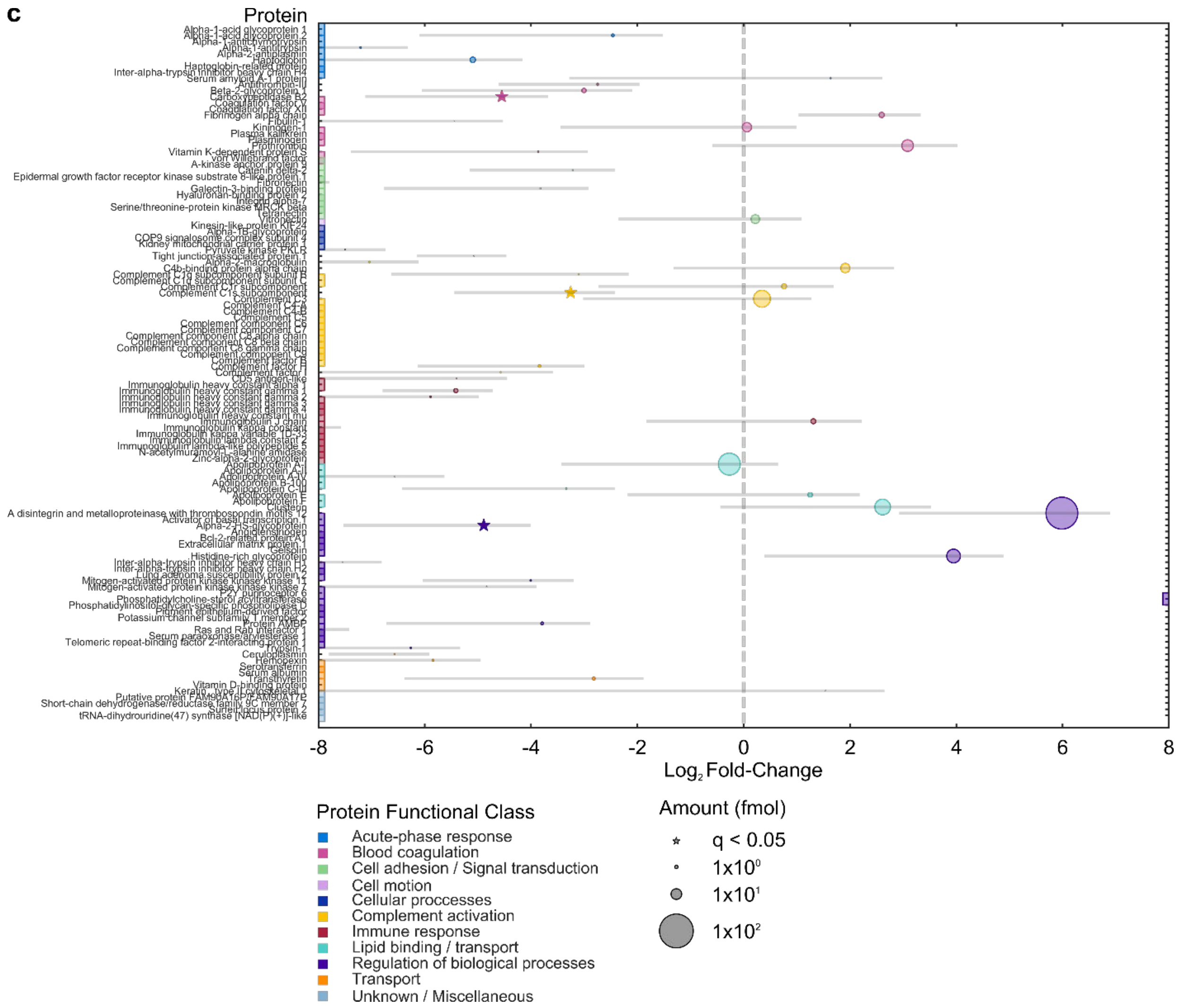


**Figure S4. Blood plasma protein corona composition determined by proteomic mass spectrometry, full results.** Protein corona formed from blood plasma on **(a)** PNPs, **(b)** (GT)$_{15}$-SWCNTs, and **(c)** (GT)$_6$-SWCNTs. Proteins are grouped by functional class according to PANTHER.[2] Fold-change is in comparison to the native biofluid alone. Circle size corresponds to the protein abundance in femtomolar. Colored boxes at x-axis limits indicate no protein detected in either corona ($x < 2^{-6}$ or $2^{-8}$) or biofluid ($x > 2^8$). Biofluid is run in technical triplicate and biofluid/nanoparticle corona samples in experimental triplicate. Error bars indicate standard error of fold-change between experimental replicates (N = 3).

**Table S1. Top 20 most abundant proteins identified by proteomic mass spectrometry in plasma alone and in $(GT)_{15}$-SWCNT and $(GT)_6$-SWCNT coronas.**

| | Plasma alone | $(GT)_{15}$-SWCNTs + plasma | $(GT)_6$-SWCNTs + plasma |
|---|---|---|---|
| 1 | Serum albumin | Clusterin | A disintegrin and metalloproteinase with thrombospondin motifs 12 |
| 2 | Haptoglobin | Histidine-rich glycoprotein | Apolipoprotein A-I |
| 3 | Ig kappa constant | Apolipoprotein A-I | Complement C3 |
| 4 | Ig heavy constant gamma | Complement C3 | Clusterin |
| 5 | Serotransferrin | Haptoglobin | Histidine-rich glycoprotein |
| 6 | Apolipoprotein A-I | A disintegrin and metalloproteinase with thrombospondin motifs 12 | Prothrombin |
| 7 | Complement C4 | Complement C1r subcomponent | Kininogen-1 |
| 8 | Telomeric repeat-binding factor 2-interacting protein | Vitronectin | C4b-binding protein alpha chain |
| 9 | Alpha-1-antitrypsin | Kininogen-1 | Vitronectin |
| 10 | Alpha-2-HS-glycoprotein | Prothrombin | Haptoglobin |
| 11 | Apolipoprotein A-II | C4b-binding protein alpha chain | Fibrinogen alpha chain |
| 12 | Ig heavy constant alpha 1 | Complement factor H | Ig J chain |
| 13 | Integrin alpha-7 | Fibrinogen alpha chain | Complement C1r subcomponent |
| 14 | Alpha-2-macroglobulin | Protein AMBP | Apolipoprotein E |
| 15 | Complement C3 | Beta-2-glycoprotein 1 | Beta-2-glycoprotein 1 |
| 16 | Complement C5 | Apolipoprotein E | Ig heavy constant gamma 1 |
| 17 | Hemopexin | Complement C1q subcomponent subunit B | Alpha-2-HS-glycoprotein |
| 18 | Alpha-1-acid glycoprotein 1 | Ig heavy constant gamma 1 | Transthyretin |
| 19 | Ig heavy constant mu | Ig J chain | Protein AMBP |
| 20 | Beta-2-glycoprotein 1 | Galectin-3-binding protein | Alpha-1-acid glycoprotein 2 |

### Proteomic mass spectrometry (LC-MS/MS) data interpretation

Prior to LC-MS/MS analysis, all samples were normalized on a total protein mass basis (where normalizing on a total molar basis is experimentally not feasible due to the complexity of biofluid samples). Consequently, the reported abundance of each protein species *i*, $b_i$, is the ratio of mole number of protein *i*, $n_i$, to the total protein mass:

$$b_i = \frac{n_i}{\sum_j n_j MW_j}$$

where $MW_j$ is the molecular weight of each protein species *j*. LC-MS/MS data is then expressed as the fold change $\varepsilon_i$ between the abundance of protein species *i* in the corona on the nanoparticle surface (phase *s*) to that in the bulk biofluid (phase *f*):

$$\varepsilon_i = \frac{b_i^s}{b_i^f} = \left(\frac{n_i^s}{n_i^f}\right)\left(\frac{\sum_j n_j^f MW_j}{\sum_j n_j^s MW_j}\right)$$

Here, the second term in parentheses is equal to 1 because all samples have the same total protein mass. Therefore, the reported fold change is the molar abundance ratio of a particular protein in the corona phase to that in the bulk biofluid phase.

**Extended discussion on plasma protein corona constituents identified by proteomic mass spectrometry**

Fibrinogen is a large rod-like multimeric protein with alpha, beta, and gamma subunits. Although identified on 2D PAGE, fibrinogen beta and gamma chains were absent from not only the nanoparticle-biofluid LC-MS/MS results, but also the native plasma samples. Additionally, there was no improvement using elution with SDS and purification with the S-trap mini column. However, the fibrinogen alpha chain was present and enriched from plasma. Based on the reproducible involvement of fibrinogen in the corona from 2D PAGE results and representation of the alpha chain in LC-MS/MS results, fibrinogen was concluded to bind to $(GT)_{15}$-SWCNTs.

For complement proteins, the particular binding mode to the nanoparticle surface results in drastically dissimilar outcomes: intact binding of complement proteins may promote nanoparticle clearance from circulation or, conversely, locally deplete complement recognition elements to effectively mask the nanoparticle's presence and thus prolong circulation *in vivo*.[3,4] SWCNTs have been previously shown to activate complement via the classical pathway,[3,5] and these findings are in agreement with our experimental result of significant complement protein adsorption. The ssDNA-SWCNT surface may present an array of adsorbed corona proteins that the complement system deems as "foreign", thus activating complement systems and leading the coated SWCNT to act as an adjuvant that increases immune response. Complement proteins may bind either directly to the SWCNT surface (as was found for complement component 1q, or C1q, on double-walled carbon nanotubes),[3] or interact with other plasma proteins adsorbed on the SWCNT (where, for example, C1q binds to immunoglobulins, IgG and IgM, or fibronectin). Due to the relatively low levels of immunoglobulins and fibronectin found in the $(GT)_{15}$-SWCNT corona, we conjecture that the former, direct binding mechanism is more likely, at least in the case of C1q (the 17$^{th}$ most abundant protein on $(GT)_{15}$-SWCNTs in plasma). Unfortunately, even a low degree of C1q binding can activate the complement system, as there are several amplification steps. Yet, if the SWCNT serves to either locally sequester proteins that initiate complement activation (such as complement C3, 4$^{th}$ most abundant) or the corona contains down-regulators in their native state (such as complement factor H, 12$^{th}$ most abundant), this could in turn bypass recognition and complement activation.

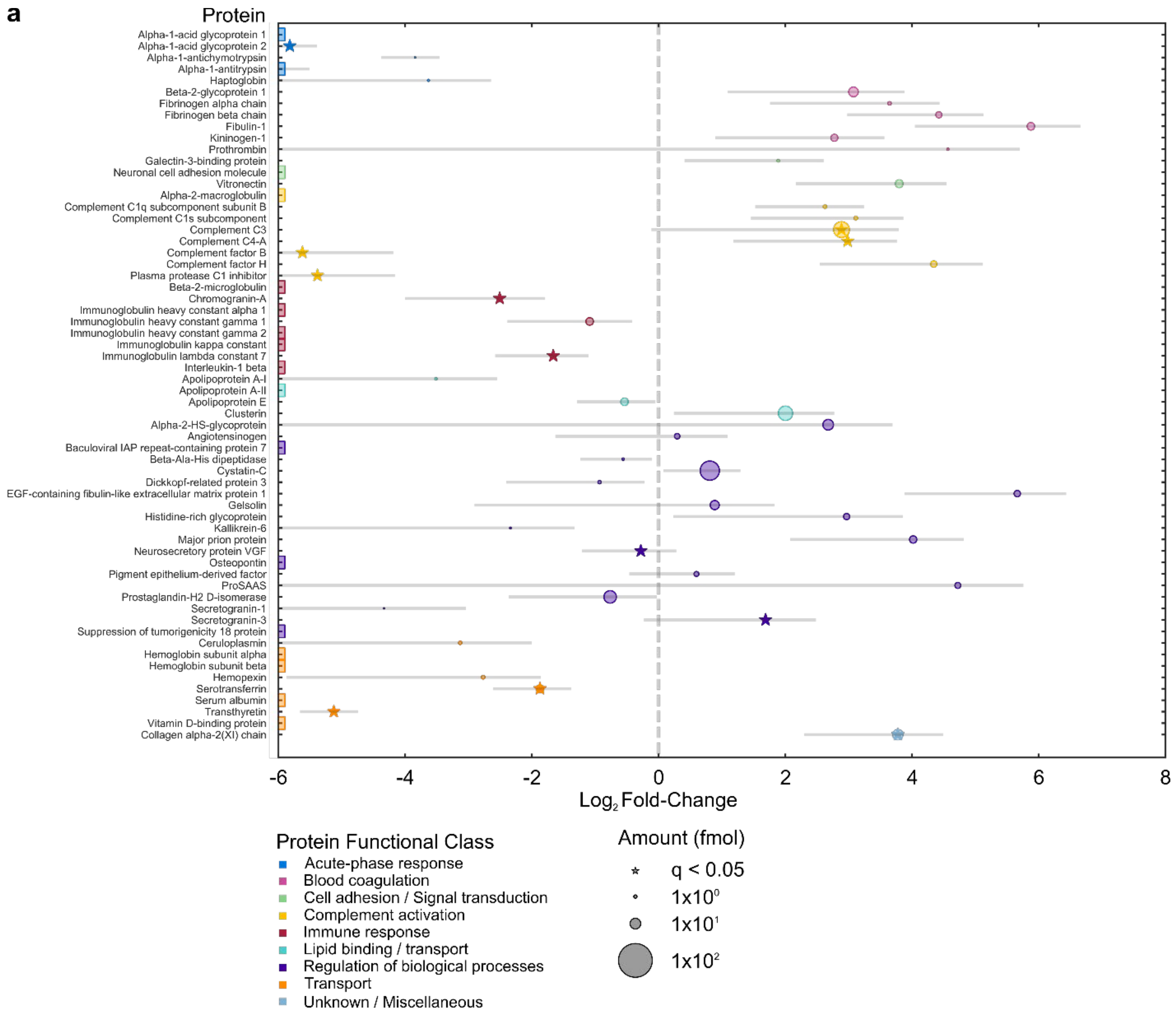
a
Protein
Alpha-1-acid glycoprotein 1
Alpha-1-acid glycoprotein 2
Alpha-1-antichymotrypsin
Alpha-1-antitrypsin
Haptoglobin
Beta-2-glycoprotein 1
Fibrinogen alpha chain
Fibrinogen beta chain
Fibulin-1
Kininogen-1
Prothrombin
Galectin-3-binding protein
Neuronal cell adhesion molecule
Vitronectin
Alpha-2-macroglobulin
Complement C1q subcomponent subunit B
Complement C1s subcomponent
Complement C3
Complement C4-A
Complement factor B
Complement factor H
Plasma protease C1 inhibitor
Beta-2-microglobulin
Chromogranin-A
Immunoglobulin heavy constant alpha 1
Immunoglobulin heavy constant gamma 1
Immunoglobulin heavy constant gamma 2
Immunoglobulin kappa constant
Immunoglobulin lambda constant 7
Interleukin-1 beta
Apolipoprotein A-I
Apolipoprotein A-II
Apolipoprotein E
Clusterin
Alpha-2-HS-glycoprotein
Angiotensinogen
Baculoviral IAP repeat-containing protein 7
Beta-Ala-His dipeptidase
Cystatin-C
Dickkopf-related protein 3
EGF-containing fibulin-like extracellular matrix protein 1
Gelsolin
Histidine-rich glycoprotein
Kallikrein-6
Major prion protein
Neurosecretory protein VGF
Osteopontin
Pigment epithelium-derived factor
ProSAAS
Prostaglandin-H2 D-isomerase
Secretogranin-1
Secretogranin-3
Suppression of tumorigenicity 18 protein
Ceruloplasmin
Hemoglobin subunit alpha
Hemoglobin subunit beta
Hemopexin
Serotransferrin
Serum albumin
Transthyretin
Vitamin D-binding protein
Collagen alpha-2(XI) chain
-6
-4
-2
0
2
4
6
8
Log2 Fold-Change
Protein Functional Class
Acute-phase response
Blood coagulation
Cell adhesion / Signal transduction
Complement activation
Immune response
Lipid binding / transport
Regulation of biological processes
Transport
Unknown / Miscellaneous
Amount (fmol)
q < 0.05
1x10^0
1x10^1
1x10^2

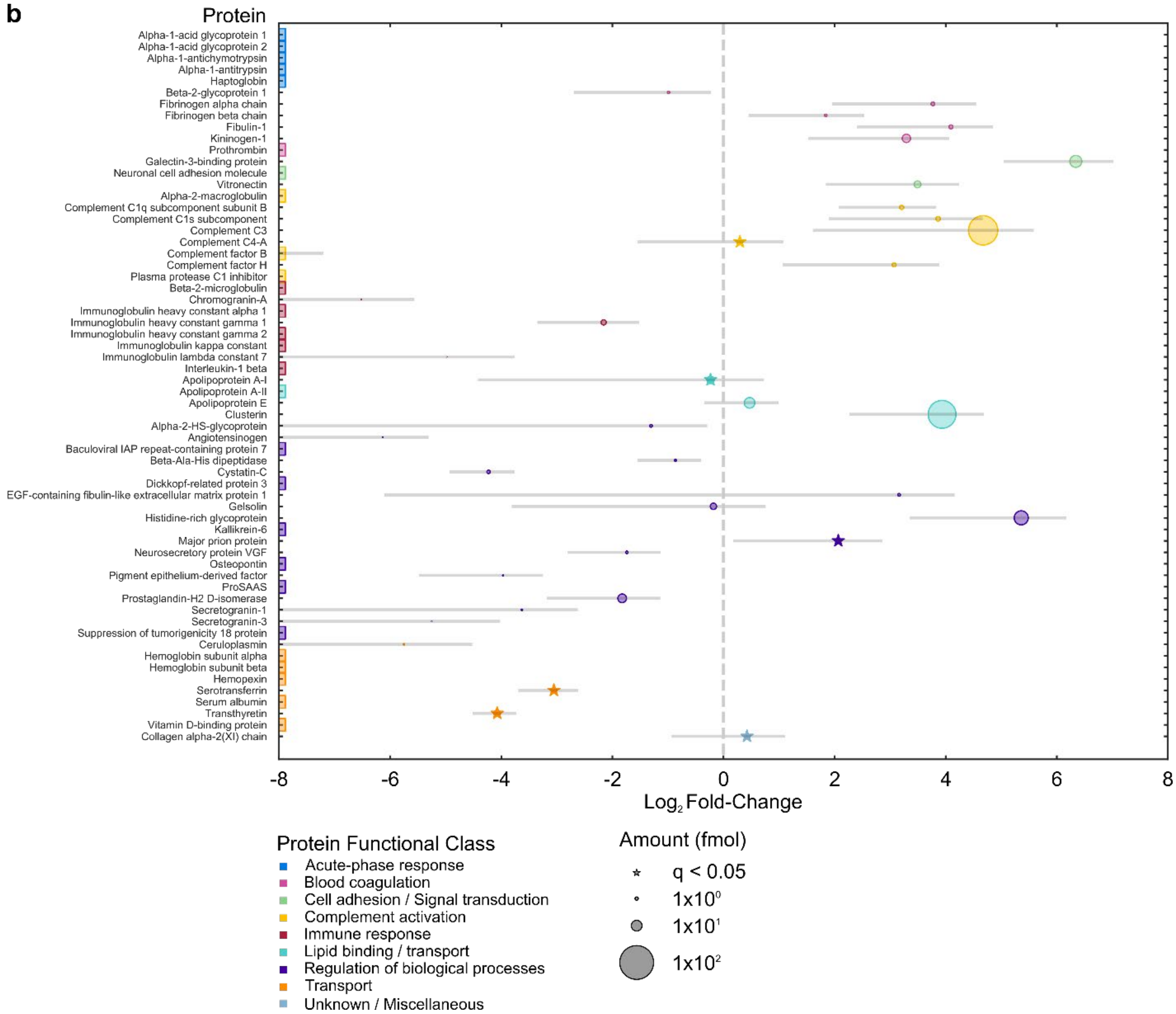


**Figure S5. Cerebrospinal fluid (CSF) protein corona composition determined by proteomic mass spectrometry, full results.** Protein corona formed from CSF on **(a)** PNPs and **(b)** $(GT)_{15}$-SWCNTs. Proteins are grouped by functional class according to PANTHER.[2] Fold-change is in comparison to the native biofluid alone. Circle size corresponds to the protein abundance in femtomolar. Colored boxes at x-axis limits indicate no protein detected in either corona ($x < 2^{-6}$ or $2^{-8}$) or biofluid ($x > 2^{8}$). Biofluid and biofluid/nanoparticle corona samples are run in technical triplicate. Error bars indicate standard deviation of fold-change between technical replicates (N = 3).

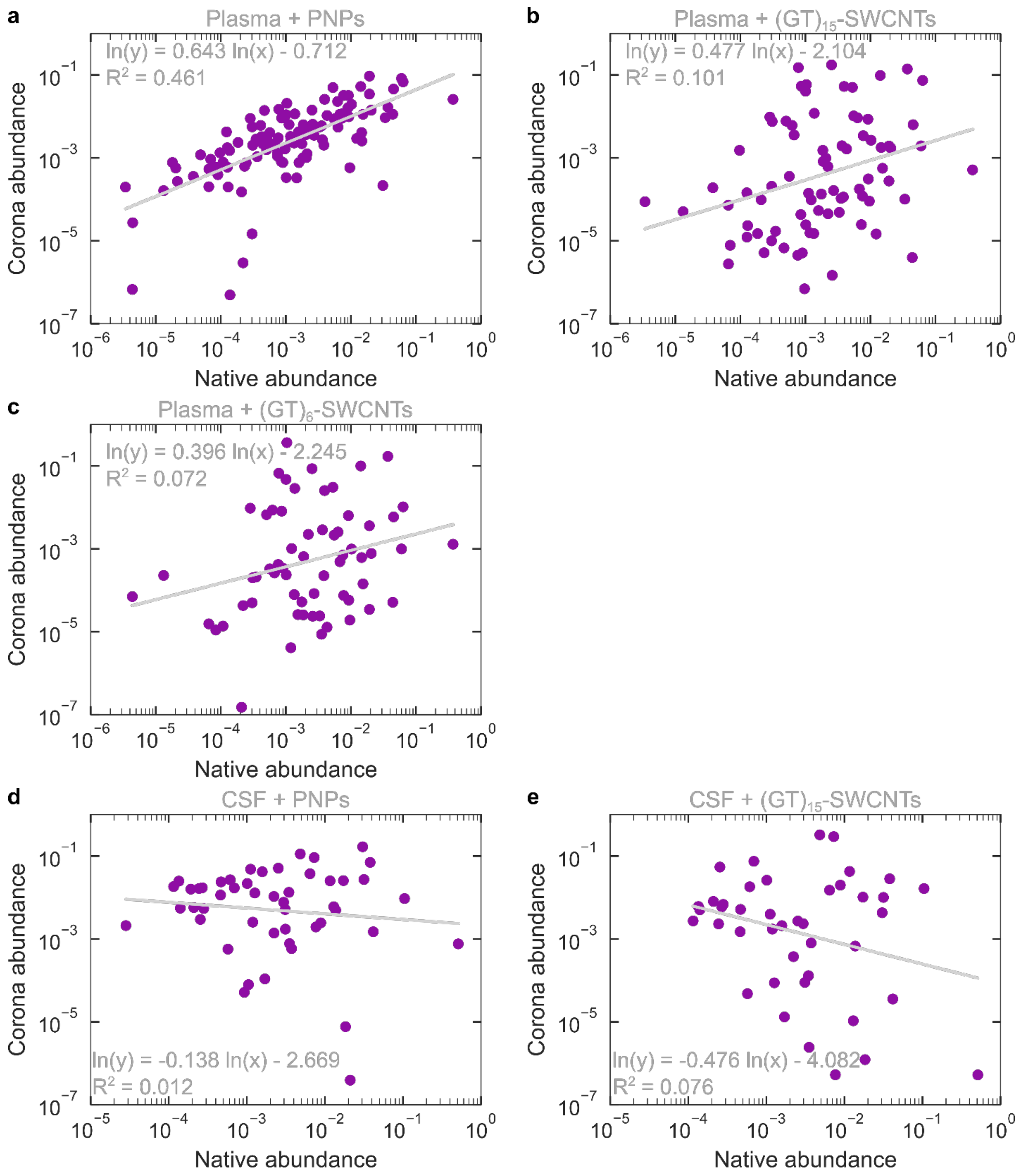


**Figure S6. Scaling of protein abundance in corona vs. in native biofluid.** Protein mole fraction of plasma proteins in corona of **(a)** PNPs, **(b)** $(GT)_{15}$-SWCNTs, and **(c)** $(GT)_6$-SWCNTs, vs. protein mole fraction of plasma proteins in native biofluid. Corona abundance scaling is approximately linear for plasma proteins on PNPs ($R^2$ = 0.461) vs. highly scattered for $(GT)_{15}$-SWCNTs ($R^2$ = 0.101) and $(GT)_6$-SWCNTs ($R^2$ = 0.072). Protein mole fraction of CSF proteins in corona of **(d)** PNPs and **(e)** $(GT)_{15}$-SWCNTs vs. protein mole fraction of CSF proteins in native biofluid. Corona abundance displays a weak negative correlation with native abundance for CSF proteins on both PNPs ($R^2$ = 0.012) and $(GT)_{15}$-SWCNTs ($R^2$ = 0.076). All mole fractions are on a solvent-free basis. Note that proteins with zero corona abundance are excluded from the analysis for clarity, but the same conclusions hold when included.

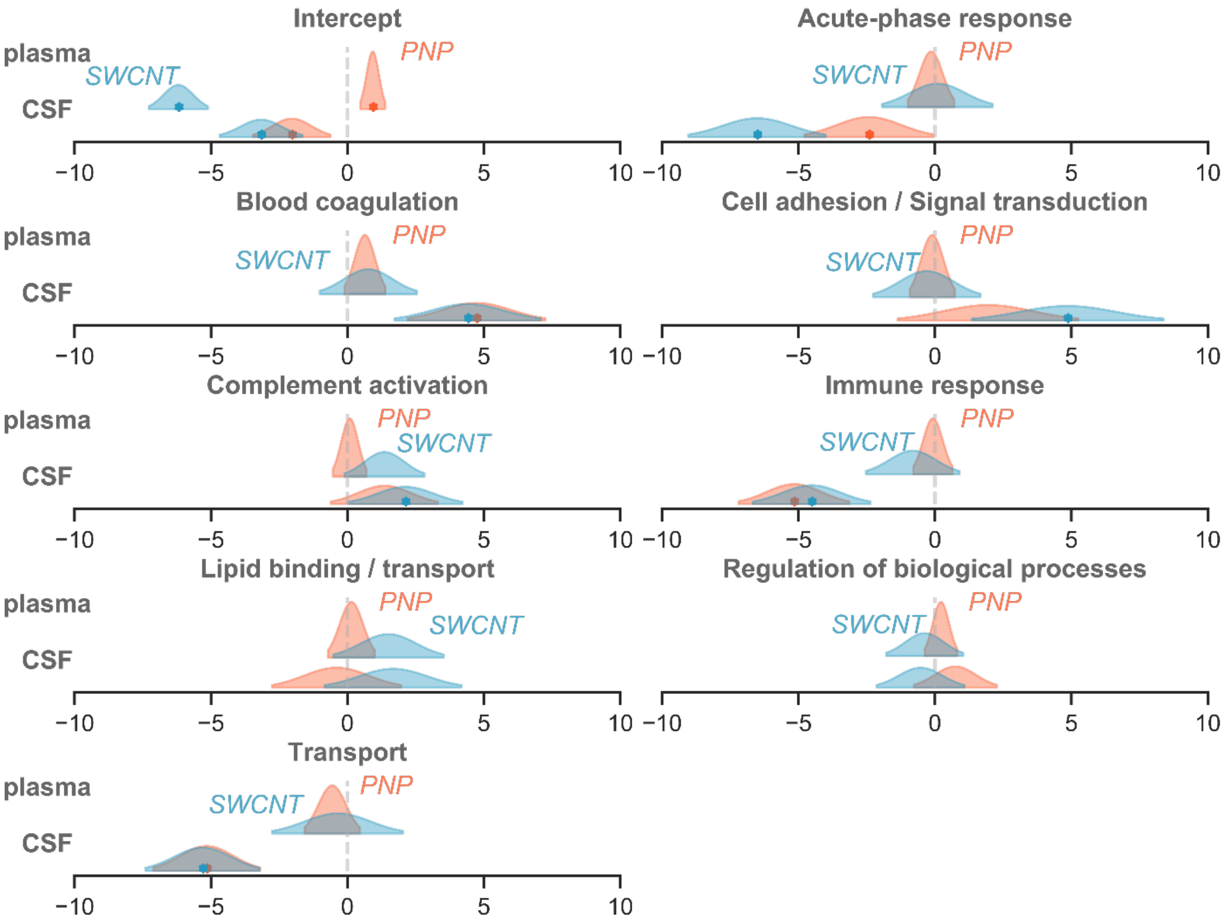


**Figure S7. Distribution for protein class regression coefficients in each nanoparticle-biofluid pairing.** Stars indicate false-discovery-rate adjusted p-values<0.1.

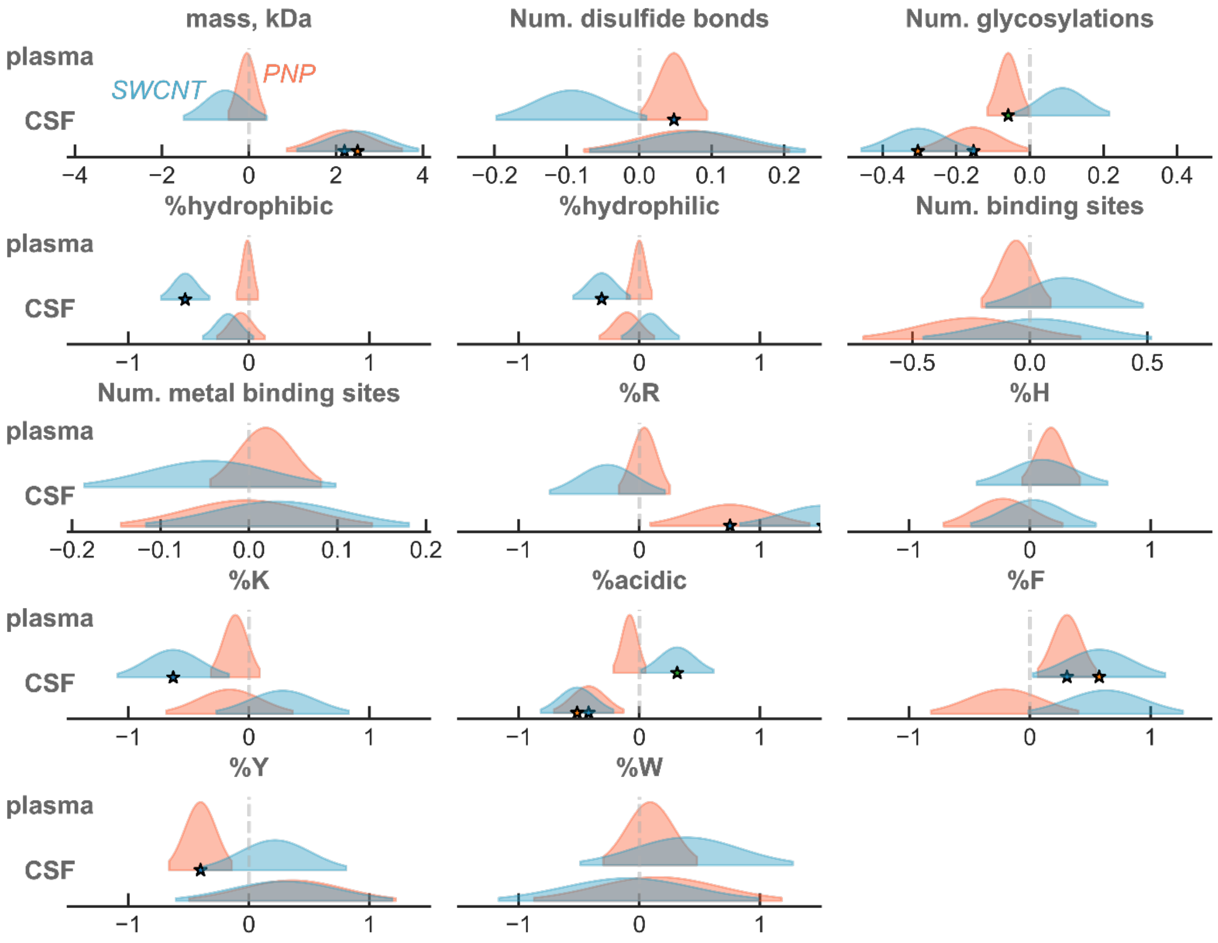


**Figure S8. Distribution for microscale regression coefficients in each nanoparticle-biofluid pairing**. Stars indicate false-discovery-rate adjusted p-values<0.1.

#### Connecting linear regression model to thermodynamics

The ideal solution chemical equilibrium constant $K_P$ of a protein in bulk solution $P$ adsorbing onto a nanoparticle into the protein corona $P^*$ is equal to the ratio of the surface concentration of the protein on the nanoparticle $\Gamma_P$ to the concentration of the protein in solution $c_P$: [6]

$$K_P = \frac{\Gamma_P}{c_P}$$

The logarithm of the chemical equilibrium constant is related to $\Delta G_P^o$, the change in standard state Gibbs free energy for a protein adsorbing from solution to the nanoparticle surface,[6]

$$\ln K_\mathrm{P} = -\frac{\Delta G_P^o}{RT}$$

where $R$ is the ideal gas constant and $T$ is temperature. The molar abundances of $P$ measured by LC-MS/MS from the native biofluid, $n_P^f$, and eluted from the nanoparticle surface, $n_P^s$, are related to $c_P$ and $\Gamma_P$ according to:

$$c_P = \frac{n_P^f f_P^f}{V}$$
$$\Gamma_P = \frac{n_P^s f_P^s}{S}$$

where $V$ is the volume and $S$ is the total surface area of the nanoparticle in the nanoparticle-biofluid solution. $f_P^f$ and $f_P^s$ are the fraction of moles of $P$ that enter the LC-MS/MS relative to amount in the fluid and on the nanoparticle surface, respectively. Dilution and steps in the pull-down assay cause $f_P^f$ and $f_P^s$ to vary from unity. This analysis neglects changes to the solution protein concentration due to corona formation. Rearrangement puts the LC-MS/MS measured log-fold change, $\ln(n_P^s/n_P^f)$, in terms of dilution factors and free energy changes:

$$\ln\left(\frac{n_P^s}{n_P^f}\right) = -\ln\left(\frac{V f_P^s}{S f_P^f}\right) - \frac{\Delta G_P^o}{RT}$$

In comparison, linear regression of the log-fold change gives:

$$\ln\left(\frac{n_P^s}{n_P^f}\right) = \beta_0 + \sum_{i=1} \beta_i X_{i,P} + \epsilon_P$$

where $\beta_i$ is a regression coefficient corresponding to $X_{i,P}$, the $i$th independent variable of protein $P$, and $\epsilon_P$ is the disturbance term that accounts for any factors other than $X_{i,P}$ controlling the fold changes.[7] Because LC-MS/MS sample preparation should not impact proteins differently, $f_P^s$ and $f_P^f$ are the same for all proteins for a set nanoparticle-biofluid system. Consequently, we can relate the chemical and statistical parameters:

$$\beta_0 = -\ln\left(\frac{V f_P^s}{S f_P^f}\right)$$
$$\sum_{i=1} \beta_i X_{i,P} + \epsilon_P = -\frac{\Delta G_P^o}{RT}$$

The regression coefficients $\beta_i$, therefore, relate the protein properties $X_i$ to the Gibbs free energy change of proteins binding to the nanoparticle $\Delta G_P^o$.

**Table S2. Protein class regression results for each nanoparticle-biofluid pairing.**

| | PNP + plasma | | | $(GT)_{15}$-SWCNT + plasma | | |
|---|---|---|---|---|---|---|
| | R-squared | Adjusted R-squared | | R-squared | Adjusted R-squared | |
| | 0.26 | 0.24 | | 0.1 | 0.13 | |
| | Parameter | Standard Error | Adjusted p-values | Parameter | Standard Error | Adjusted p-values |
| **Intercept** | 0.9277 | 0.2270 | 0.0001 | -6.1942 | 0.5380 | 0.0000 |
| **Acute-phase response** | -0.1393 | 0.4271 | 0.7445 | 0.0916 | 1.0124 | 0.9279 |
| **Blood coagulation** | 0.6394 | 0.3749 | 0.0891 | 0.7639 | 0.8887 | 0.3907 |
| **Cell adhesion / Signal transduction** | -0.0981 | 0.4133 | 0.8125 | -0.3003 | 0.9797 | 0.7594 |
| **Complement activation** | 0.0854 | 0.3086 | 0.7822 | 1.3564 | 0.7316 | 0.0647 |
| **Immune response** | -0.0727 | 0.3620 | 0.8410 | -0.8149 | 0.8580 | 0.3430 |
| **Lipid binding / transport** | 0.1467 | 0.4271 | 0.7315 | 1.5017 | 1.0124 | 0.1390 |
| **Regulation of biological processes** | 0.2203 | 0.2985 | 0.4609 | -0.3749 | 0.7075 | 0.5966 |
| **Transport** | -0.5563 | 0.5047 | 0.2713 | -0.3584 | 1.1965 | 0.7647 |
| **Sample 1** | 2.8743 | 0.3253 | 0.0000 | 4.0556 | 0.7711 | 0.0000 |
| **Sample 2** | -0.0358 | 0.3146 | 0.9095 | 0.3037 | 0.7457 | 0.6841 |

| | PNP + CSF | | | $(GT)_{15}$-SWCNT + CSF | | |
|---|---|---|---|---|---|---|
| | R-squared | Adjusted R-squared | | R-squared | Adjusted R-squared | |
| | 0.32 | 0.28 | | 0.35 | 0.31 | |
| | Parameter | Standard Error | Adjusted p-values | Parameter | Standard Error | Adjusted p-values |
| **Intercept** | -2.0347 | 0.7108 | 0.0048 | -3.1619 | 0.7551 | 0.0000 |
| **Acute-phase response** | -2.4143 | 1.1830 | 0.0429 | -6.5197 | 1.2566 | 0.0000 |
| **Blood coagulation** | 4.7294 | 1.2588 | 0.0002 | 4.4095 | 1.3371 | 0.0012 |
| **Cell adhesion / Signal transduction** | 1.9484 | 1.6552 | 0.2409 | 4.8610 | 1.7583 | 0.0064 |
| **Complement activation** | 1.3603 | 0.9806 | 0.1673 | 2.1252 | 1.0417 | 0.0430 |
| **Immune response** | -5.1599 | 1.0141 | 0.0000 | -4.5191 | 1.0772 | 0.0000 |
| **Lipid binding / transport** | -0.3967 | 1.1830 | 0.7378 | 1.6716 | 1.2566 | 0.1853 |
| **Regulation of biological processes** | 0.7459 | 0.7590 | 0.3272 | -0.5229 | 0.8062 | 0.5175 |
| **Transport** | -5.1597 | 0.9806 | 0.0000 | -5.3045 | 1.0417 | 0.0000 |
| **Sample 1** | 0.9624 | 0.8879 | 0.2800 | -0.0962 | 0.9431 | 0.9188 |
| **Sample 2** | 0.8264 | 0.8915 | 0.3553 | -0.0073 | 0.9470 | 0.9939 |

**Table S3. Microscale regression results for each nanoparticle-biofluid pairing.**

| | PNP + plasma | | | $(GT)_{15}$-SWCNT + plasma | | |
|---|---|---|---|---|---|---|
| | R-squared | Adjusted R-squared | | R-squared | Adjusted R-squared | |
| | 0.32 | 0.29 | | 0.27 | 0.23 | |
| | **Parameter** | **Standard Error** | **Adjusted p-values** | **Parameter** | **Standard Error** | **Adjusted p-values** |
| **Intercept** | 2.7690 | 3.6300 | 0.4462 | 19.6116 | 8.2339 | 0.0179 |
| **Sample 1** | 2.8759 | 0.3147 | 0.0000 | 4.0662 | 0.7137 | 0.0000 |
| **Sample 2** | -0.0427 | 0.3036 | 0.8881 | 0.3186 | 0.6886 | 0.6439 |
| **Mass** | -0.0475 | 0.2109 | 0.8219 | -0.5446 | 0.4785 | 0.2560 |
| **% hydrophobic residues (nonaromatic)** | -0.0126 | 0.0442 | 0.7759 | -0.5286 | 0.1003 | 0.0000 |
| **% hydrophilic residues** | -0.0008 | 0.0520 | 0.9877 | -0.3122 | 0.1179 | 0.0085 |
| **% acidic residues** | -0.0806 | 0.0662 | 0.2243 | 0.3131 | 0.1502 | 0.0379 |
| **% arginine** | 0.0409 | 0.1051 | 0.6975 | -0.2643 | 0.2385 | 0.2687 |
| **% histidine** | 0.1762 | 0.1197 | 0.1422 | 0.1028 | 0.2716 | 0.7053 |
| **% lysine** | -0.1128 | 0.1015 | 0.2672 | -0.6276 | 0.2302 | 0.0068 |
| **% phenylalanine** | 0.3077 | 0.1209 | 0.0114 | 0.5747 | 0.2743 | 0.0370 |
| **% tyrosine** | -0.4013 | 0.1297 | 0.0022 | 0.2165 | 0.2942 | 0.4624 |
| **% tryptophan** | 0.0893 | 0.1940 | 0.6455 | 0.3912 | 0.4400 | 0.3747 |
| **Number of disulfide bonds** | 0.0477 | 0.0229 | 0.0379 | -0.0938 | 0.0519 | 0.0719 |
| **Number of glycosylated sites** | -0.0588 | 0.0280 | 0.0370 | 0.0885 | 0.0636 | 0.1654 |
| **Number of ligand binding sites** | -0.0584 | 0.0736 | 0.4276 | 0.1473 | 0.1668 | 0.3779 |
| **Number of metal binding sites** | 0.0190 | 0.0313 | 0.5452 | -0.0442 | 0.0710 | 0.5340 |

| | PNP + CSF | | | $(GT)_{15}$-SWCNT + CSF | | |
|---|---|---|---|---|---|---|
| | R-squared | Adjusted R-squared | | R-squared | Adjusted R-squared | |
| | 0.35 | 0.29 | | 0.4 | 0.34 | |
| | **Parameter** | **Standard Error** | **Adjusted p-values** | **Parameter** | **Standard Error** | **Adjusted p-values** |
| **Intercept** | -18.8198 | 9.7672 | 0.0558 | -32.4292 | 10.2373 | 0.0018 |
| **Sample 1** | 0.7932 | 0.8845 | 0.3712 | -0.4235 | 0.9271 | 0.6485 |
| **Sample 2** | 0.7215 | 0.8883 | 0.4179 | -0.2622 | 0.9311 | 0.7786 |
| **Mass** | 2.1956 | 0.6640 | 0.0012 | 2.4973 | 0.6960 | 0.0004 |
| **% hydrophobic residues (nonaromatic)** | -0.0670 | 0.0987 | 0.4981 | -0.1726 | 0.1034 | 0.0972 |
| **% hydrophilic residues** | -0.1034 | 0.1133 | 0.3628 | 0.0913 | 0.1187 | 0.4429 |
| **% acidic residues** | -0.4209 | 0.1441 | 0.0040 | -0.5162 | 0.1511 | 0.0008 |
| **% arginine** | 0.7511 | 0.3295 | 0.0240 | 1.5253 | 0.3453 | 0.0000 |
| **% histidine** | -0.2217 | 0.2464 | 0.3696 | 0.0293 | 0.2583 | 0.9097 |
| **% lysine** | -0.1607 | 0.2616 | 0.5398 | 0.2798 | 0.2742 | 0.3090 |
| **% phenylalanine** | -0.2088 | 0.3052 | 0.4950 | 0.6294 | 0.3199 | 0.0509 |
| **% tyrosine** | 0.3624 | 0.4282 | 0.3986 | 0.2910 | 0.4488 | 0.5177 |

| | | | | | | |
|---|---|---|---|---|---|---|
| **% tryptophan** | 0.1523 | 0.5128 | 0.7669 | -0.0914 | 0.5375 | 0.8653 |
| **Number of disulfide bonds** | 0.0653 | 0.0708 | 0.3577 | 0.0801 | 0.0742 | 0.2819 |
| **Number of glycosylated sites** | -0.1529 | 0.0733 | 0.0387 | -0.3039 | 0.0769 | 0.0001 |
| **Number of ligand binding sites** | -0.2462 | 0.2313 | 0.2888 | 0.0309 | 0.2424 | 0.8987 |
| **Number of metal binding sites** | -0.0028 | 0.0709 | 0.9685 | 0.0323 | 0.0744 | 0.6647 |

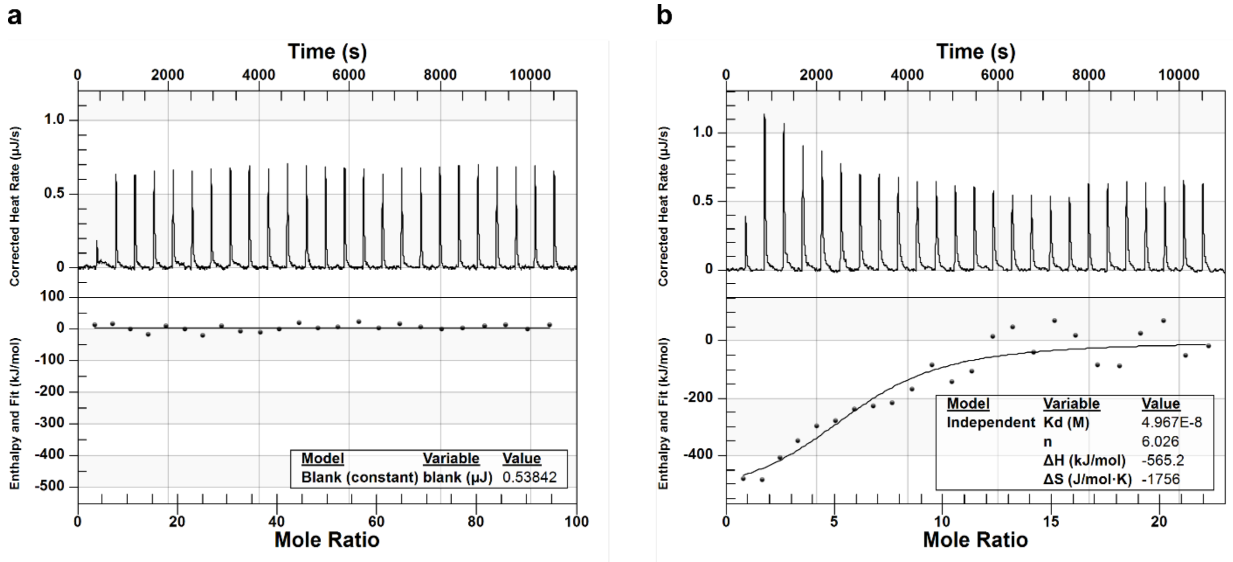


**Figure S9. Protein corona thermodynamics assessed with ITC for binding of key proteins to $(GT)_{15}$-SWCNTs.** Isothermal titration calorimetry (ITC) is employed to determine binding thermodynamics of **(a)** albumin and **(b)** fibrinogen to $(GT)_{15}$-SWCNTs. Albumin does not bind to $(GT)_{15}$-SWCNTs within experimentally accessible limits of this instrument, whereas fibrinogen does, in agreement with the corona compositional analyses from proteomic mass spectrometry and gel electrophoresis.

### Isothermal titration calorimetry (ITC) methods

ITC measurements were performed with a NanoITC (TA Instruments). Prior to each experiment, samples and buffer were degassed for 10 min and the reference cell was filled with fresh Milli-Q water. Equilibration time was set to 1 h before the experiment start and the initial and final baselines were collected for 300 s. For each experiment, 1.2 g/L protein in 0.1 M PBS was titrated from the syringe (250 µL total volume) into 0.1 g/L $(GT)_{15}$-SWCNTs in 0.1 M PBS in the cell (1 mL total volume) under constant stirring (250 rpm) at 25 °C. 10 µL of protein titrant was injected into the nanoparticle solution in the cell every 7 min, with a total of 24 injections. By standard practice, every run was initiated with a 5 µL injection to ensure no artifacts due to bubbles and was removed from analysis. All protein-nanoparticle binding experiments were accompanied by three heat-of-dilution control experiments: (1) protein injected into buffer, (2) buffer injected into nanoparticles, and (3) buffer injected into buffer (where buffer is 0.1 M PBS). Heat of binding of protein to nanoparticles was then calculated as: (heat from titration of protein into nanoparticles) – (1) – (2) + (3). Data processing was completed with NanoAnalyze software (TA Instruments). Baseline correction was done using the auto-fit routine. An independent binding model was applied to fit the fibrinogen data set, suitable to model weak nonspecific interactions such as those present in the system under study,[8] and a blank (constant) model was applied to fit the albumin data set.

Protein and nanoparticle concentrations and ITC setup parameters were varied in attempt of obtaining binding curves for both proteins to $(GT)_{15}$-SWCNTs. However, for albumin this was not possible within the ITC instrument's operational range, therefore albumin was concluded to not bind to $(GT)_{15}$-SWCNTs.

**Extended discussion on isothermal titration calorimetry (ITC)**

ITC was employed to extract relative binding parameters of protein-nanoparticle association. ITC was performed at constant pressure such that the heat absorbed or released is equivalent to the change in enthalpy ($\Delta H^o$) upon binding. The binding curve can be fit to determine the equilibrium dissociation constant ($K_d$) and molar binding stoichiometry (n). This enables subsequent calculation of changes in standard state Gibbs free energy ($\Delta G^o$) and entropy ($\Delta S^o$) as follows:

$$\Delta G^o = RT\ln K_d = \Delta H^o - T\Delta S^o$$

where $R$ is the ideal gas constant and $T$ is temperature. The optimized run parameters to measure heats of binding for this system require relatively high protein and nanoparticle concentrations. At these concentrations, addition of fibrinogen causes visible sample aggregation, presumably due to polymer bridging interactions of proteins adsorbed on one nanoparticle interacting with another nanoparticle. One of the key assumptions of ITC is that the system is equilibrated during each titration step. Yet, aggregation is a kinetically controlled, non-equilibrium process. As the key assumption is not held, these binding values are actually the convolution of protein binding to individual SWCNTs, fibrinogen binding to aggregated SWCNTs, and SWCNTs aggregating. We can compensate for this limitation in data processing by applying the Lumry-Eyring model,[9] in which an equilibrium reaction is coupled to a self-association reaction (i.e. aggregation), and the heats measured are separated out accordingly. This encompasses subtracting out baseline aggregation heats and arriving at an apparent binding heat. Therefore, the thermodynamic parameters are reported with consideration of these higher order processes taking place simultaneously. A further note is that baseline drift/shift were observed during these ITC experiments involving $(GT)_{15}$-SWCNTs. These changes in baseline often indicate slow non-equilibrium processes in action, further confirming the presence of aggregation. In conclusion, ITC is not a suitable methodology to study nanoparticle-protein corona formation for all systems, and these limitations must be considered during experimental design and reporting of results.

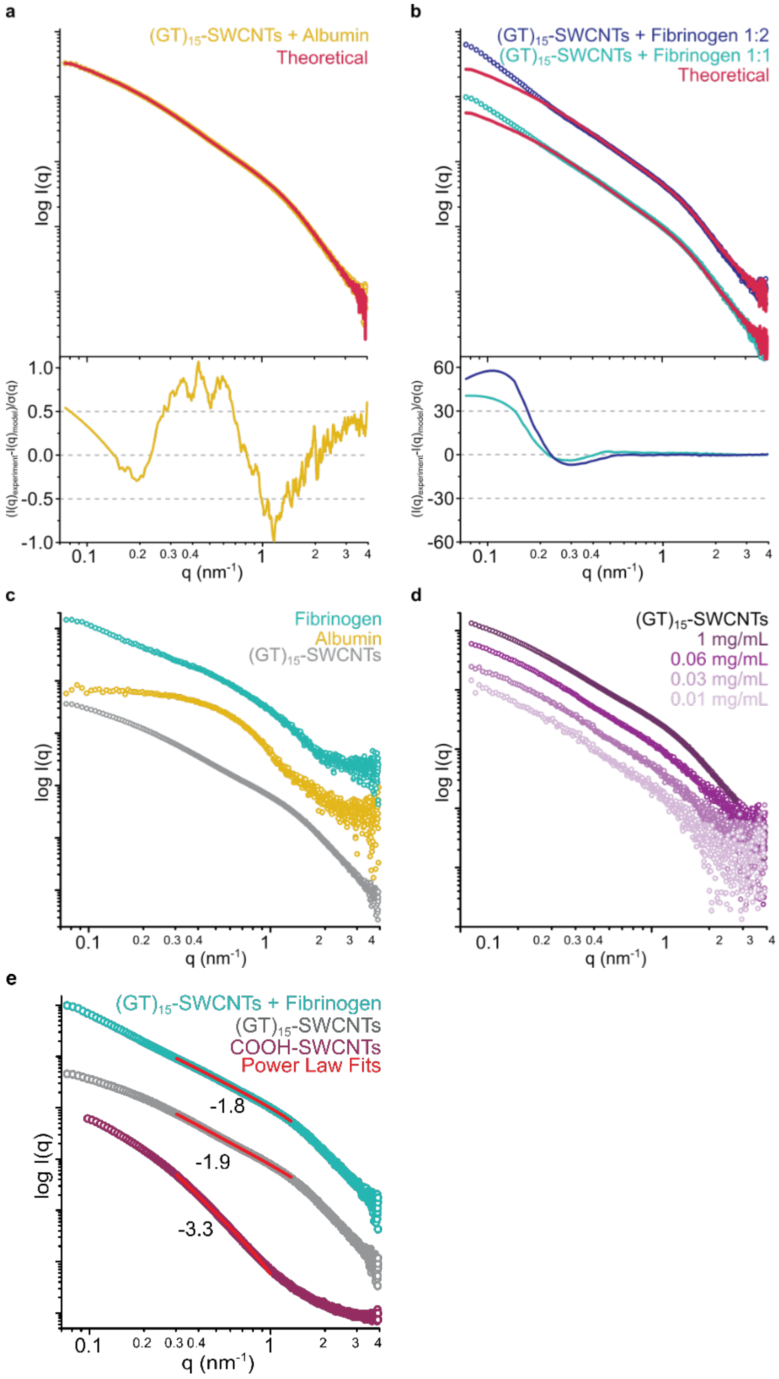


**Figure S10. Protein corona structure assessed with SAXS for binding of key proteins to (GT)15-SWCNTs.** The linear combination of respective standard curves from panel c in red and fit-residuals below, fit against the curves produced by the potential complexes of (GT)15-SWCNTs with **(a)** albumin or **(b)** fibrinogen, at two different ratios of (GT)15-SWCNTs to fibrinogen (1:1 is 0.5 g/L final concentrations of (GT)15-SWCNTs and fibrinogen; 1:2 is 0.25 g/L (GT)15-SWCNTs and 0.5 g/L fibrinogen). **(c)** Experimental SAXS profiles for standards of albumin, fibrinogen, and (GT)15-SWCNTs alone, at identical concentrations to the mixing experiments. **(d)** SAXS profiles for concentration series of (GT)15-SWCNTs. **(e)** SAXS profiles fit to show power law dependencies in the Porod regions, including the COOH-SWCNT control without surface-adsorbed ssDNA.

**Table S4. SAXS mass fractal modeling parameters.**

| Sample | Radius (nm) | Fractal Dimension | Cutoff Length (nm) |
|---|---|---|---|
| $(GT)_{15}$-SWCNTs + Fibrinogen | 1.05 ± 0.003 | 1.77 | 103.34 ± 9.70 |
| $(GT)_{15}$-SWCNTs + Albumin | 1.05 ± 0.003 | 1.90 | 10.60 ± 0.05 |
| $(GT)_{15}$-SWCNTs | 1.01 ± 0.002 | 1.89 | 10.91 ± 0.04 |

**Extended modeling details and discussion on small-angle x-ray scattering (SAXS)**

Scattering profiles were fit using the mass fractal model. These fits were complimented by determining the power-law dependency of the Porod region and were both calculated using the SasView software package (www.sasview.org). Scattering intensity as a function of scattering vector $I(q)$ calculations for the mass fractal modeling (**Figure 7**) was done as follows:[10]

$$I(q) = \text{scale} * P(q)S(q) + \text{background}$$

$$P(q) = F(qR)^2$$

$$F(\text{x}) = \frac{3[\sin(x) - x\cos(x)]}{x^3}$$

$$S(q) = \frac{\Gamma(Dm-1)\zeta^{Dm-1}}{[1+(q\zeta)2]^{\frac{Dm-1}{2}}}\frac{\sin[(Dm-1)\tan^{-1}(q\zeta)]}{q}$$

$$\text{scale} = \text{scale factor} * N\left(\frac{4}{3}\pi R^3\right)^2(\rho_{\text{particle}} - \rho_{\text{solvent}})^2$$

where $R$ is the radius of the building block, $Dm$ is the mass fractal dimension, $\zeta$ is the cut-off length, $N$ is number of scatters, $\rho_{solvent}$ is the scattering length density of the solvent, and $\rho_{particle}$ is the scattering length density of particles. $Dm$ relates the mass ($m$) to the radius as $m \sim R^{Dm}$ and is analogous to $I(q) \sim q^{-p}$ from the power-law calculations (with power-law exponent $p$), where $Dm = p$ when $q\zeta >> 1$.

The power-law dependencies were determined by fitting the experimental SAXS profiles (**Figure 7**), where $0.3 \leq q \leq 1$ nm$^{-1}$ with the following:[11]

$$I(q) = \text{scale} * q^{-p} + \text{background}$$

These power-law dependencies were then used to confirm the calculated $Dm$ from the mass fractal model fits, as $Dm$ is analogous to $p$ in the power-law calculations. Mass fractal modeling parameters are tabulated in **Table S4**.

Control SAXS profiles of albumin, fibrinogen, and $(GT)_{15}$-SWCNTs alone were collected at identical concentrations to those of the mixing experiments (**Figure S10c**). Data was collected at elevated concentrations (0.5 mg/mL both protein and $(GT)_{15}$-SWCNTs) to enhance SAXS signal, however, a concentration series was also performed for $(GT)_{15}$-SWCNTs to ensure that the scattering profiles do not deviate when under more biologically relevant conditions down to 0.01 mg/mL (**Figure S10d**).[12] As another control, carboxylic acid functionalized SWCNTs (COOH-SWCNTs) were also examined via power-law scattering obtaining $p \sim 3.3$ (**Figure S10e**). This fit suggests that without ssDNA functionalization, COOH-SWCNTs form roughly spherical aggregates better modeled as a uniform density as opposed to a polymeric mass fractal. Thus, it may be inferred that ssDNA provides some semblance of order to the fine molecular structure of the system and should be the subject of further investigation.

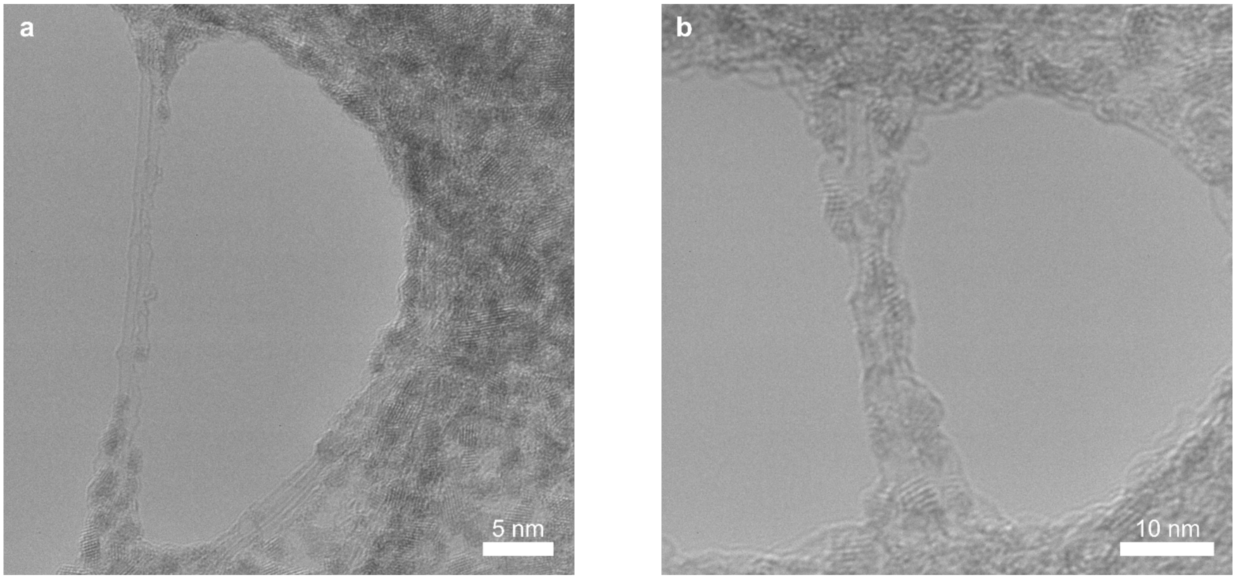


**Figure S11. Protein corona morphology visualized by TEM for adsorption of plasma proteins to $(GT)_{15}$-SWCNTs.** Transmission electron microscopy (TEM) of **(a)** plasma protein corona and **(b)** fibrinogen corona on $(GT)_{15}$-SWCNTs.

### Transmission electron microscopy (TEM) methods

Holey carbon-coated grids (EMS Electron Microscopy Science) were surface-treated by glow discharge to make the support hydrophilic. Samples of $(GT)_{15}$-SWCNTs with fibrinogen or plasma were negatively stained with 1% uranyl acetate solution. For the $(GT)_{15}$-SWCNTs alone sample, no negative staining was done. 5 µl of 10 mg/L solution was drop-cast onto the grid. FEI ThemIS 60-300 STEM/TEM (National Center of Electron Microscopy, Molecular Foundry) with acceleration voltage of 60kV was used to acquire TEM images by video recording. A low acceleration voltage was chosen to minimize sample damage and increase sample contrast.

**Table S5. Purchased biofluid and protein specifications.**

| Protein | Manufacturer | Lot # | Source | Form |
|---|---|---|---|---|
| Blood plasma | Innovative Research Inc. | #23791 | Pooled normal human plasma | Biofluid |
| Cerebrospinal fluid | Lee Biosolutions | #07C5126 | Pooled normal human CSF, from remnant lumbar puncture | Biofluid |
| Albumin | Sigma-Aldrich | #SLBZ2785 | Human plasma | Lyophilized |
| Alpha-2-HS glycoprotein | Biovision Inc. | #4C08L75480 | Human plasma | Lyophilized |
| Apolipoprotein A-I | Alfa Aesar | #927J17A | Human plasma | 1mg/ml in 10mM ammonium bicarbonate buffer, pH 7.4 |
| Clusterin | R&D Systems | NEV1519031 | Mouse myeloma cell line, NS0-derived human; Asp23-Arg227 (beta) & Ser228-Glu449 (alpha) with a C-terminal 6-His tag | Lyophilized |
| Complement C3 | Mybiosource Inc. | #N30/20170 | Human plasma | 5 mg/mL |
| Fibrinogen | Millipore Sigma | #3169957 | Human plasma | Lyophilized |
| Immunoglobulin G | Lee Biosolutions | #06B2334 | Human plasma | Lyophilized |
